# Restriction Enzyme activity under Force determined with autonomous DNA Origami Force Clamps via Gel Electrophoresis

**DOI:** 10.64898/2026.11.13.765095

**Authors:** Arthur Ermatov, Michael Matthies, Petr Šulc, Tim Liedl

## Abstract

Investigating force-depending behaviour in biological systems is a challenging task, with most techniques requiring complex experimental equipment while also often being limited in scalability. We designed a facile force spectroscopy technique using DNA origami-based force clamps that apply forces up to ∼40 pN. We validated the approach by using type II endonucleases that have been previously studied quantitatively by single-molecule measurements. Using ensemble measurements based on ubiquitously used agarose gel electrophoresis, we qualitatively observed force-dependent behaviour for EcoRV and force-independent behaviour for BamHI, replicating previous data for these enzymes. Interestingly, we also observed force-dependent behaviour for EcoRI, whose restriction behaviour has been previously described as force-independent. Our experimental design opens a path to force spectroscopy based on bottom-up techniques and ensemble measurements, allowing to collect statistical data in large quantities with readily available techniques in many wet labs.

## Introduction

Forces acting on biomolecular systems play a crucial role in their behaviour. RNA polymerase, for example, stalls at forces above ∼14 pN [1], kinesin motors at forces below 8 pN [2] and myosin V already at 5 pN [3]. Directed forces in cells also play a key role in shaping the exoskeleton and arranging cell organelles [4]. Several methods exist to investigate force-dependent behaviour of cells, proteins and nucleic acids on the nanometer- and on the nano-to femtonewton scale. When investigating force-dependent behaviour of biological systems, single-molecule tools are often the method of choice as they they can reveal molecular sub-populations, short-lived or transient states, rare events and complex energy landscapes. Optical or magnetic tweezers are, for example, being used routinely to study single-molecular behaviour of DNA strands [5, 6]. Proteins are often probed through pulling them apart with an atomic force microscope (AFM) [7]. Generally, by recording the pulling force and the extension of the molecule of interest these methods allow force spectroscopy over a wide range of forces (femtonewton - hundreds of nanonewton) [8].

However, preparing individual DNA strands or proteins for single-molecule measurements is a tedious, multi-step process. To acquire optical single-molecule data, specialised microscope setups such as total internal reflection or confocal microscopes have to be used that are not readily available in all labs. Further, several measurements have to be performed to obtain reproducible, statistically relevant results; ensemble measurements require a lot of experimental effort, if possible at all.

Other techniques allow higher throughputs when performing measurements. Magnetic tweezers allow the observation of multiple molecules tethered to magnetic beads in a single experiment at low magnification, and thus a wide field of view [9]. In an alternative approach, called centrifugal force microscopy, arrays of tethered molecules are exposed to centrifugal forces and and monitored with optical methods [10, 11]; pulling forces can also be applied by virtue of acoustic [12, 13] and flow forces [14]. Compared with AFM or optical tweezers, many more molecules can be observed here in parallel.

Yet another approach relies on bottom-up fabrication and makes use of the molecular control offered by DNA origami. After demonstrating that single-stranded DNA (ssDNA) can be tensed by rigid DNA origami struts in so-called tensegrity (a portmanteau of tension and integrity) configurations [15], our group implemented a DNA origami force clamp that is able to exert forces up to 12 pN [16]. These force clamps were read out using single-molecule Fluorescence Resonance Energy Transfer (FRET) both on substrates and in bulk. By immobilising the DNA origami structures on glass substrates, FRET time traces could be recorded, at the expense of limited data acquisition band width. In bulk, a confocal single-molecule setup with alternating laser excitation (ALEX) allowed recording tens of thousands of events in 15 minutes. The same force clamp was later applied to study RNA Polymerase III pre-initiation complex stability [17]. Using a different DNA origami-based force spectrometer design, Funke et al. derived the distance-dependent energy landscape for nucleosome pair interactions [18] and Darcy et al. used a similar design to apply forces between 10 and 20 pN [19]. Zhou et al. employed DNA origami force clamps as molecular tools to study mechanosensitive proteins on a biochemical scale up to 13 pN [20].

All these methods still have in common that they rely on sensitive fluorescent microscopes to record the individual FRET events. In this study, we investigated the performance of restriction enzymes using widely available agarose gel-electrophoresis as the readout mechanism.

Next to their fundamental roles in biology, restriction enzymes are an important tool in molecular biology. Initially discovered as a mechanism to defend bacteria from viruses by cleaving viral DNA, they have been studied and characterised extensively [21, 22]. By recognising specific double-stranded DNA (dsDNA) sequences mostly between 4 and 8 base pairs long, they allow for precise DNA cleavage at sites of interest for many different applications such as gene editing and synthetic biology, making them indispensable in today’s microbiology labs. Today, they are readily commercially available, and a large variety of restriction enzymes are used in molecular biology [23].

In this work, we demonstrate a force tool based on DNA origami that applies tunable forces up to 40 pN. We show a proof-of-concept application of this device and perform ensemble force measurements with a variety of DNA endonucleases. As read-out we use agarose gel electrophoresis. To verify and fine-tune the forces applied on the DNA strands, we developed a computational method based on MD simulations, adapting and generalising the theoretical approach provided by various models for force-dependent behaviour of single- and double-stranded DNA.

## Results

Resembling earlier published work [16], we modified our U-shaped force clamp design consisting of a 100 nm long “base” and two “tower” regions. Between these towers, single-stranded scaffold DNA of controllable lengths is tensed, thus acting as an entropic spring. To increase mechanical stability and reduce the torque applied to the towers, we placed the ssDNA region as close to the base as possible. The scaffold section between the towers also contained the multiple cloning site (MCS, Supporting Information for sequence details). By adding oligonucloetides that were complementary to the restriction sites under investigation, a double-stranded region under tension was generated (see Supporting Information for design and experimental details).

To easily implement the different force clamp variants, each with their designed applied force, only a small portion of the staple strands – so called “force staples” – had to be replaced in each design. We were thus able to modify the number of single-stranded nucleotides between the two tower regions of the DNA origami, thereby changing the length of the scaffold region and hence the forces acting on the the dsDNA region of interest (Figure 1a) [15].

**Figure 1.**
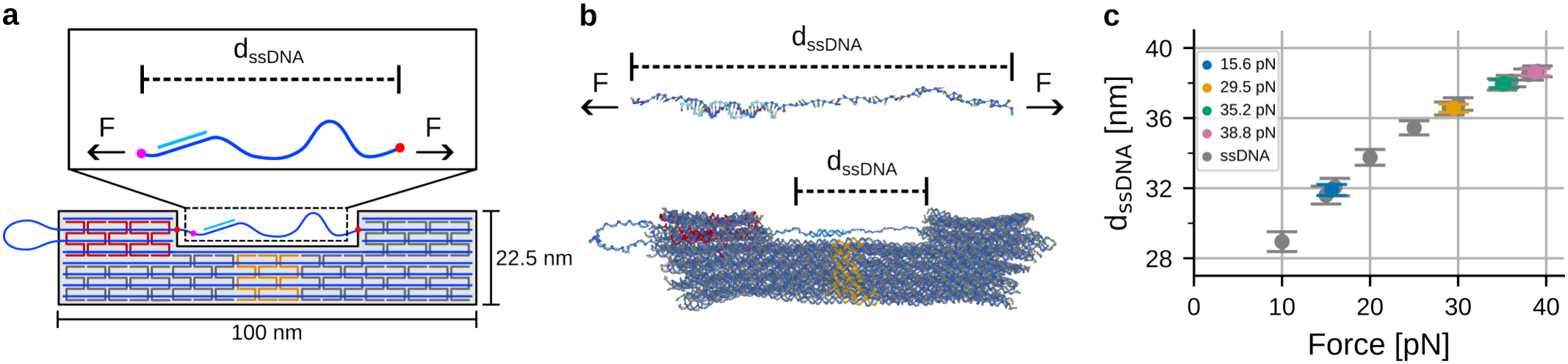
Design of DNA origami force clamp structure. **a**, Scheme of U-shaped DNA origami design. A ssDNA region (in between the red dots) that is part of the scaffold acts as an entropic spring. The entropic spring carries a double-stranded region (bright blue) that is used for force validation experiments. The number of ssDNA nucleotides determines the strain applied to the ssDNA region. A region conserved over all force clamp desings (between red and purple dot) is used to simulate the extension d_(_dsDNA) under different applied forces. **b**, oxDNA simulations of ssDNA region set under pre-defined strain (top) and of the entire DNA origami structure (bottom). Base markers have been omitted in the origami image for clarity. Note that the scale of the two simulation images differs. **c**, Simulated end-to-end distances of ssDNA region (grey) for specific forces compared to corresponding end-to-end distances of the conserved region inside different DNA origami designs.

To apply accurate predefined strains, several models and approaches were considered. The mFJC model provides a sufficient approximation of single-stranded DNA being strained under force [24], while double-stranded DNA is best described by the worm-like chain (WLC) model [25] [26]. While the theoretical limit for our force clamp is given by force-induced melting of double-stranded DNA in the origami structure, the real limit of our design is significantly lower since DNA origami structures are based on short dsDNA regions interconnected via

Holliday junctions instead of longer dsDNA regions [27]. In addition, the mFJC and WLC models do not take into account the flexibility of the structure or any modifications such as a dsDNA subregion within the ssDNA region. Further, the models behave ill-conditioned for higher forces: individual nucleotides (un)binding result in changes of several pN in high force regimes (*±* 5 pN for 40 pN designed force and higher, see Figure S1 and Table S1).

Based on the standard modified freely-jointed chain (mFJC) model, we first designed our structures to exert forces on a double-stranded region in the range of 10, 30, 50 and 60 pN (Figure S1). We then approximated the applied forces empirically by comparing the end-to-end distance of a conserved region within the DNA origami structure comprising the strained ssDNA and the dsDNA reporter region to the same region being pulled along its principal axis with pre-defined forces (Figure 1a) using oxDNA molecular dynamics (MD) simulations [28, 29].

Applying different simulated forces onto the isolated conserved region (Figure 1b, top) allowed us to generate calibration data that maps specific ssDNA extension distances to the applied forces that in turn would be required to achieve these distances (Figure 1c, grey circles). We then compared these distances to the distances obtained from oxDNA simulations of the conserved regions in the DNA origami structures we designed (Figure 1b, bottom) to estimate the applied force on this region within the structures (Figure 1c, coloured circles). The two distance values obtained from the conserved region simulation that were closest to the respective DNA origami data defined the upper and lower bounds for the applied force inside the origami structure; mapping the origami data to the linear interpolation between these two boundaries resulted in the force values shown in this work.

At forces above 10 pN we observed in our previous designs [16] occasionally broken tower regions. To avoid mechanical stress during the assembly, we established a two-step annealing process: During a first temperature ramp from 65°C to 25°C over the course of approximately 37h, the central staples of the base were absent from the folding mixture (orange staples in 1a). As a result, the “spring” remained without tensions and the towers and the larger fractions of the base could assemble undisturbed (Figure S2). In the second annealing step at 37°C for 12h, now with the central staples present (Figure 2a), the structure tensed itself by incorporating these oligonucleotides (Figure 2a).

**Figure 2.**
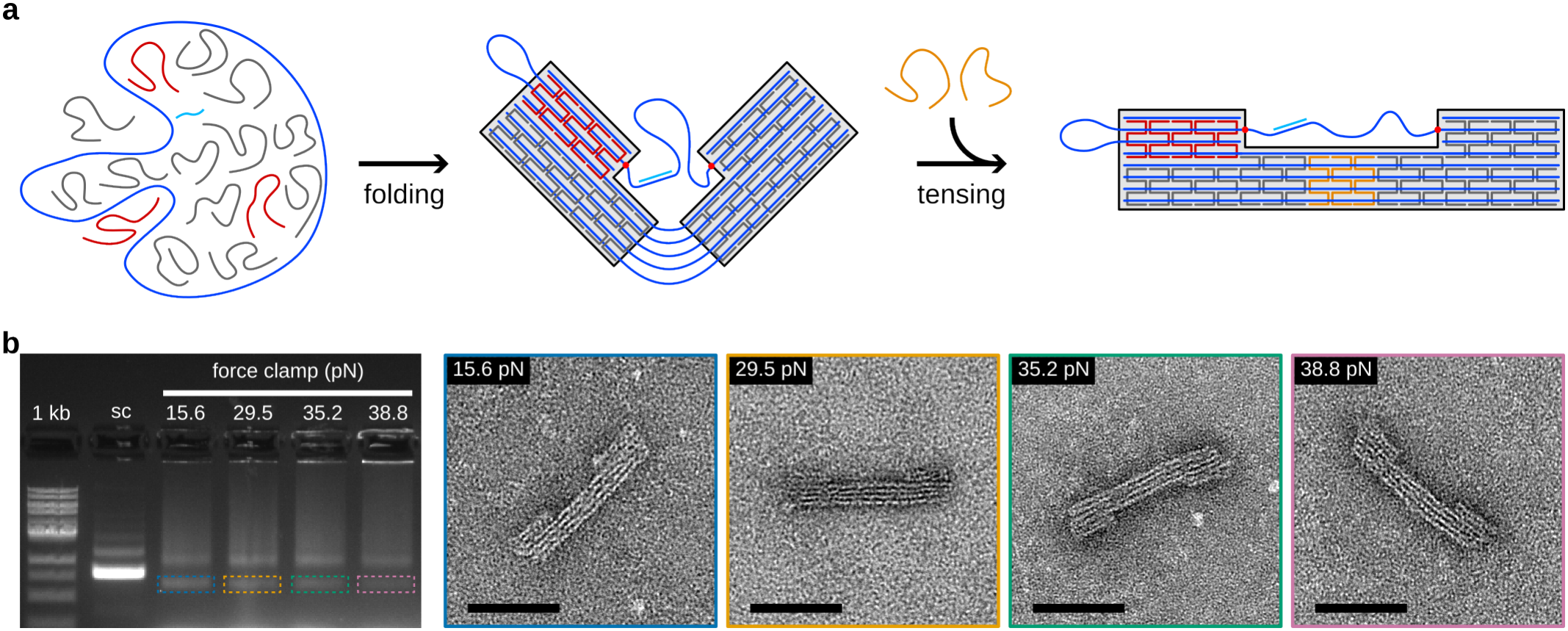
Assembly and characterization of DNA origami force clamps. a, Two-step assembly of DNA origami structures. First, the tower regions are folded leaving out the central staples (orange) of the “seam”. In a second step, the force clamps are completed and tensed by adding the seam staples. b, Characterization of structures. Left: Agarose gel electrophoresis of 1 kb ladder (1 kb), DNA origami scaffold (sc) and structures with designed forces (see labels). Gel was stained using SYBR Safe and structure samples of monomer band (coloured boxes) were extracted from gel by cutting and squeezing. Right: TEM images of force clamps with colour-coded frames (cf. left gel image), all scale bars 50 nm.

After the two-step assembly, we analyzed the different force samples with agarose gel electrophoresis and transmission electron microscopy (Figure 2b). The gel image reveals two distinct populations, represented by each two leading bands that were identified as monomers (Figure 2b) and dimers (Figure S3). Higher-order aggregates appear as smeared out intensities and material stuck in the gel pockets. With shortening ssDNA spring region and thus increasing force, the intensity of the momomer band diminishes further and further and aggregation prevails. We attribute this to the following: the center staples are designed to “close” the two open interfaces of one structure in the second folding step. However, they might as well connect the four, six, etc. interfaces of two, three, etc. structures. With shortening springs and thus increasing forces, the tensing process becomes slower and the formation of non-monomeric structures will become energetically more favourable. Another factor could involve hydrophobic effects as nucleotides get exposed to water in the tensed single-strands.

Regardless, since DNA origami is a bottom-up nanofabrication approach, this allows us to produce a large number of tensed structures simultaneously. We therefore were able to chose a technically simple and widely accessible method to perform ensemble measurements on the structures to validate our simulation data and study the influence of force on the catalytic acivity of three different restriction enzymes (Figure 3): EcoRV is a class type II endonuclease that bends the DNA duplex by 52 degrees to access and cleave its recognition site. Consequently, EcoRV has been shown to exhibit a force-dependent restriction rate in single-molecule experiments [30]. In contrast, EcoRI – while also known to twist and bend DNA – is considered to create too little bending for a force-dependent behaviour, which, to our knowledge, has so far not been interrogated at forces above 20 pN; at forces below 20 pN, no force dependency has been observed previously [31]. As a negative control enzyme, we chose BamHI which is known not to bend the DNA in the cleaving process [30]. T he force calibration simulations (Figure 1) used the scaffold sequence that included an EcoRV endonuclease recognition site within its dsDNA region of interest. We assumed the same applied forces for the structures designed to investigate EcoRI and BamHI, as long as the total number of ssDNA nucleotides and the length of the dsDNA domain were the same for all three DNA origami designs.

**Figure 3.**
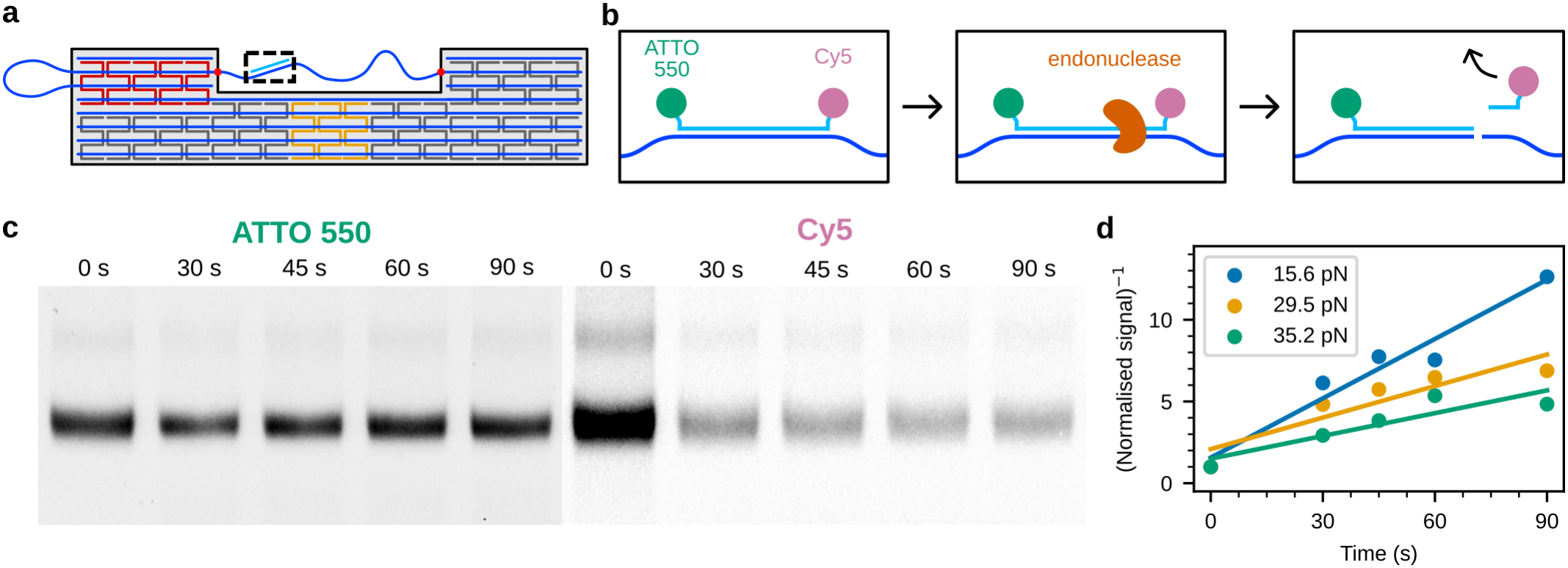
Experimental setup for force validation. **a**, Within the ssDNA region, a 28 bp long dsDNA probe region (black box) contains the restriction site and is set under tension. **b**, The probe strand carries two fluorophores (ATTO550 and Cy5) at its 5’ and 3’ ends. After cutting by the enzyme, the shorter end carrying the Cy5 dye (6 nt for EcoRV, 8 nt for EcoRI, 4 nt for BamHI) detaches from the scaffold. textbfc, Gel scanner image showing the leading band intensities for ATTO550 and Cy5. The reaction times (here with enzyme EcoRV) are indicated above each lane. **d**, Normalized enzyme activity for enzyme EcoRV under 3 forces plotted over reaction time.

For the cutting activity experiments we used a staple strand carrying Cy5 and ATTO550 at its respective strand ends that binds to the complementary sequence located on the single-stranded region that forms the entropic spring, forming a strained dsDNA probe region. The probe region encodes the recognition and ligation site of an endonuclease that binds and cuts the dsDNA. The restriction site is located in such a manner that the probe strand was cut asymmetrically, leaving one short end with Cy5 attached and one long end with ATTO550 attached. The short end carrying the Cy5 molecule then detached from the structure, leaving solely the ATTO550 molecule attached to the DNA origami scaffold strand. The reaction was then quenched at various time points after initiation using a 6x concentrate containing 90 mM ethylenediaminetetraacetic acid (EDTA) that was subsequently diluted in the reaction solution to a final concenctration of 15 mM to chelate all free Mg^2+^ ions (11 mM) available. The quenched reaction mixture was then transferred onto a Mg^2+^-free agarose gel to perform gel electrophoresis and subsequent imaging (Figure 3c). While DNA origamis normally do not maintain their structural integrity under these conditions due to reduced charge shielding by Mg^2+^ ions (Figure S4), dsDNA domains long enough to have melting temperatures above the gel running temperature – such as the probe dsDNA array carrying the fluorophores – stayed intact (Figure 3c, ATTO550 signal). As expected, the short dsDNA fragments dissociated from the gel (Figure 3c, Cy5 signal). Therefore, the Cy5 signal decreases with longer incubation times due to longer enzymatic activity and resulting restriction. A normalized signal is determined by dividing the time-dependent Cy5 signal (time-dependent) by the ATTO550 signal (time-independent) for each band, correcting absolute errors in origami concentration (see Supporting Information).

For the cutting activity experiments we added unpurified folded structures to an enzymatic reaction mixture containing a high concentration of enzyme (200 U). At such high concentrations we expect restriction activity – and not diffusion – to be the rate-limiting step. As expected, we observed constant ATTO550 signals for all forces at all incubation times within a few percent of the mean value since the DNA fragment carrying the ATTO550 molecule is long enough to stay bound the complement region on the scaffold. For all enzymes, we see a reduction in Cy5 signal for increased incubation times, with varying degrees of attenuation depending on the applied force for EcoRV and EcoRI. Based on the signal attenuation, we derived the restriction activity for every enzyme and force, respectively (see Supporting Information, Figures S5 and S6). Applying a second-order rate equation model to the normalised data gave us different restriction activities for different forces, represented by different regression slopes (Figure 3d), which we found to be a plausible model choice for enzymatic reactions with one substrate and a high enzyme-to-substrate ratio [32]. For every force, we performed a total of four (EcoRI, BamHI) or eight (EcoRV) replicates.

As expected, we see a force-dependent behaviour of EcoRV and no force-dependent behaviour of BamHI (Figure 4). Interestingly, we also observe a force-dependent behaviour for EcoRI, even though, similarly to BamHI, this enzyme has not been described as force-dependent in the past [31]. In our experiments, the force range was well above the one investigated previously. A possible explanation for the observed force dependence at higher forces could lie in the required distortion of the restriction domain of EcoRI to cut the restriction site, leading to unstacking and wedging the DNA by 55°and thus resulting in an overall kink of 12°. [21].

**Figure 4.**
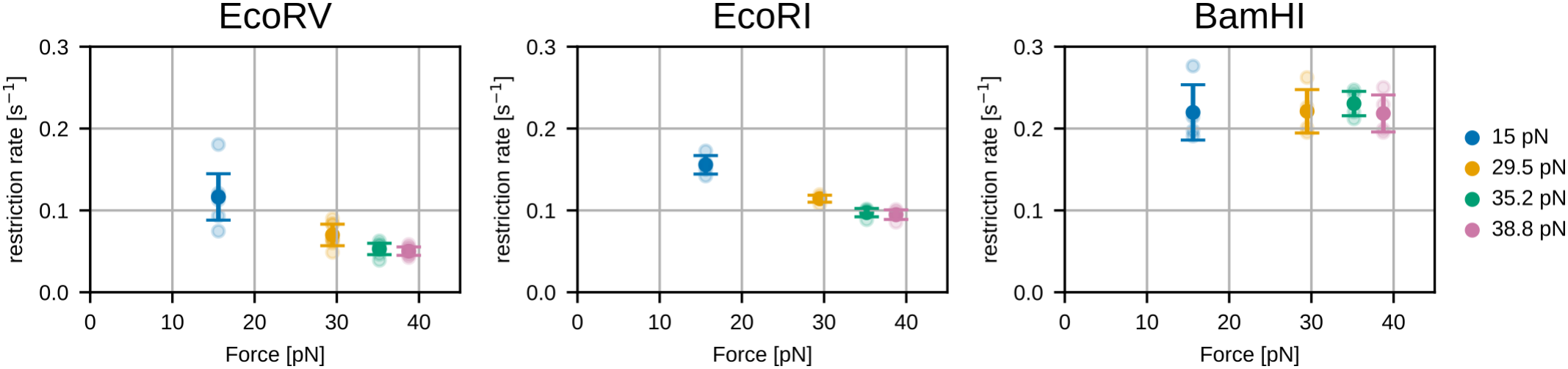
Restriction rates of different type II endonucleases. While EcoRV and EcoRI are known to bend DNA to different degrees when cutting, BamHI keeps it in a straight configuration. This is reflected in force-dependent restriction rates for EcoRV and EcoRI in contrast to force-independent restriction rates for BamHI.

## Discussion

Ensemble measurements of single-molecule kinetics using agarose gel electrophoresis are an elegant and facile way of measuring the enzymatic activity of restriction enzymes. While our experiments show the expected force-dependent enzymatic activity of EcoRV and BamHI and insights into the behavior of EcoRI when interacting with dsDNA under high forces, several limitations of this simple, bulk experiment-based methodolgy remain. Firstly, the data can only be used for a qualitative assessment of the enzymatic behaviour with respect to different applied forces since the experimental setup did not allow for any separation between monomer and dimer DNA origami structures. In particular, it is unclear what kind of force the dimers are experiencing. In principle it would be possible to purify the monomers and perform the same set of experiments on monomers only, however, this results in large variations of origami concentrations leading to ill-defined cutting conditions. In addition, the signal strength is significantly weaker due to lower origami concentrations, leading to large errors. In the experimental design chosen by us, errors in concentrations and hence signals are predominantly based on pipetting errors, which are on the order of 5% or less. For BamHI, the cutting activity appears to decline for longer incubation times (Figure S7a). We attribute this to poor signal-to-noise ratios for the low fluorescence intensities observed here as well as substrate depletion effects at longer reaction times. Consequently, we observe a better model fit when only considering data up to 30 seconds of reaction time (Figure S7b) compared to fits based on the whole dataset (Figure S7c). Importantly, all these effects appear to be independent of the applied force, thereby validating our qualitative assessment of force-independent restriction of BamHI (Figure S7c).

While we have used this technique to qualitatively assess ensemble performance, the technique itself also allows for quantitative measurements. This requires precise control over the used concentrations of the DNA nanostructures, their structural homogeneity, and an imaging setup that resolves the fluorescent signal of the gel bands to a sufficient detail.

With our initial designs intended to apply higher forces based on the mFJC model, we observed that the forces observed in the simulations were significantly smaller. We see several reasons for this. Firstly, the mFJC model is based on several parameters such as Kuhn length and the Young’s modulus of ss- and dsDNA, each based on averages of previous measurements and thus involving an error. This is of particular importance for high forces since small deviations in parameters result in large force discrepancies. Secondly, the mFJC model assumes a constant number of nucleotides for the ssDNA portion. In DNA origami structures based on dsDNA, fraying events at the ends of staple strands happen constantly based on thermal fluctuation. In large force regimes, this results in significant changes in number of nucleotides, and thus in resulting entropic spring force, similar to the error associated with deviations in other parameters of the mFJC model described earlier. Further, we observe melting of staples at the inner staples of the DNA origami adjacent to the ssDNA region. Classically, DNA origamis based on a hexagonal lattice have dsDNA regions of mostly 7 base pairs or integer multiples thereof between Holliday junctions. In our design, we chose to place a larger number of Holliday junctions in our design as we assumed that this way, we can most effectively divert mechanical strain from the structure. However, we mostly observed staple melting at the staples adjacent to the ssDNA region and no breaking of the structure. Higher forces could therefore be achieved by designing longer dsDNA regions at the bordering region to the ssDNA region, with forces over 40 pN being feasible (Figure S8).

The demonstrated method to estimate the force applied by the entropic spring inside the DNA origami structure is also capable of simulating the force applied in the ssDNA region even when modifications (e.g. protein attachment) are performed. It therefore allows for simple empirical force calibration when models might be too complex to accurately describe the experimental design.

## Acknowledgements

The authors thank Johann Moritz Weck and Eduard Unterauer for their assistance in gel imaging and Susanne Kempter for laboratory support. A.E. and T.L. acknowledge funding through the Deutsche Forschungs-gemeinschaft (DFG; German Research Foundation) through the cluster of excellence Biosystem-Design Munich (BioSysteM, EXC 3092) and the Free State of Bavaria through the ONE MUNICH Project Munich Multiscale Biofabrication. M.M. and P.Š. acknowledges funding from the European Research Council under the European Union’s Horizon 2020 research and innovation program (grant agreement number 101040035).

## Supporting information

The Supporting Infomation is available free of charge.

- Experimental and analysis protocols, TEM images of dimers and destroyed structures after gel electrophoresis, caDNAno designs of DNA origami structures, DNA sequences of scaffolds and staple strands.

## Notes

### Competing Interest Statement

The authors have declared no competing interest.

