## Supporting Information for "Restriction Enzyme activity under Force determined with autonomous DNA Origami Force Clamps via Gel Electrophoresis"

#### mFJC force model compared to MD simulation data

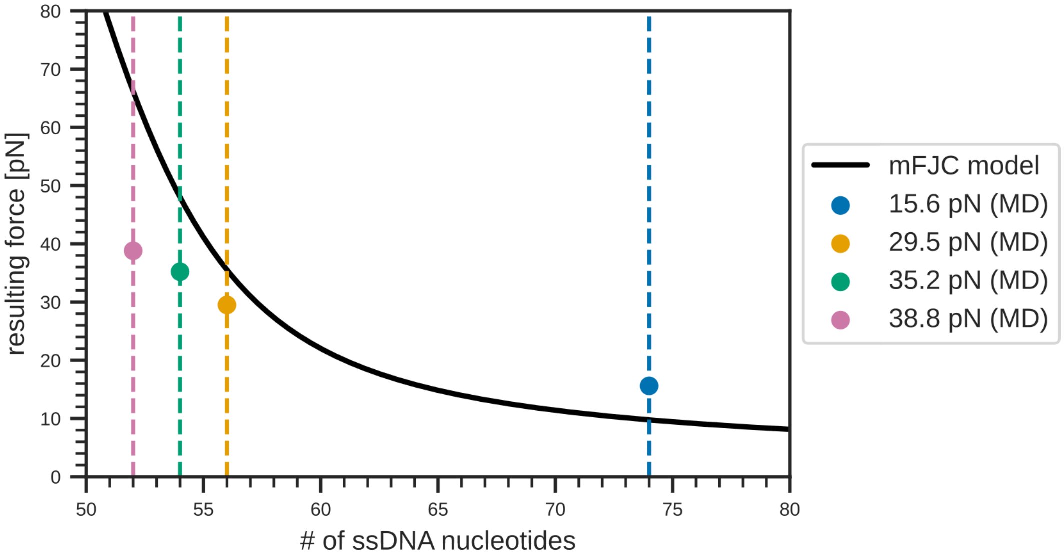


**Figure S1:** Prediction of entropic spring force in the different DNA origami structure designs. Four structures were designed with a total number of free nucleotides in the entropic spring region being 74, 56, 54, and 52 nucleotiudes respectively. The forces determined by molecular dynamics (MD) simulations (coloured) are compared to the predictions of the mFJC model for a specific number of nucleotides (black line).

| # of free nt | MD force [pN] | mFJC force [pN] | mFJC force range [pN] |
|---|---|---|---|
| 74 | 15.6 | 9.8 | 9.6 - 9.9 |
| 56 | 29.6 | 35.6 | 33.2 - 38.1 |
| 54 | 35.5 | 48.0 | 44.4 - 51.9 |
| 52 | 38.8 | 66.2 | 61.1 - 71.6 |

**Table S1:** Overview of force predictions for a specific number of free nucleotides in the entropic spring region of the DNA origami structure. The molecular dynamics (MD) simulations analysed the entropic spring region with the dsDNA fluorescent reporter region incorporated in the origami design, with the number of free nucleotides of the entropic spring being split by the reporter region. The mFJC analysis considered an entropic spring solely consisting of ssDNA.

### DNA origami assembly

#### Folding protocol

The concentrations shown in Table S2 were used for a total folding volume of 50 µl aqueous solution. The folding mix was subsequently heated according to the temperature ramp described in Table S3.

| Component | Concentration |
|---|---|
| $MgCl_2$ | 20 mM |
| Tris-EDTA buffer | 10mM/1mM |
| p8634 scaffold | 30 nM |
| Force staples | 300 nM |
| Core staples | 300 nM |
| Reporter staple | 300 nM |

**Table S2:** DNA origami folding mixture.

| Temperature [℃] | Time [min:seconds] |
|---|---|
| 65 | 01:53 |
| 64-61 | 04:45 |
| 60-59 | 20:15 |
| 58 | 42:45 |
| 57 | 65:45 |
| 56 | 88:15 |
| 55 | 110:15 |
| 54-45 | 132:45 |
| 44 | 110:15 |
| 43 | 87:45 |
| 42 | 65:15 |
| 41-39 | 42:45 |
| 38-37 | 20:15 |
| 36-30 | 10:54 |
| 29-26 | 01:53 |
| 25 | hold |

**Table S3:** Temperature ramp for DNA origami folding. Samples were held at 25℃ until further processing.

After folding, seam staples were added (volume equivalent to 300 nM in 50 µl), and the structures were incubated for 12 hours at 37℃.

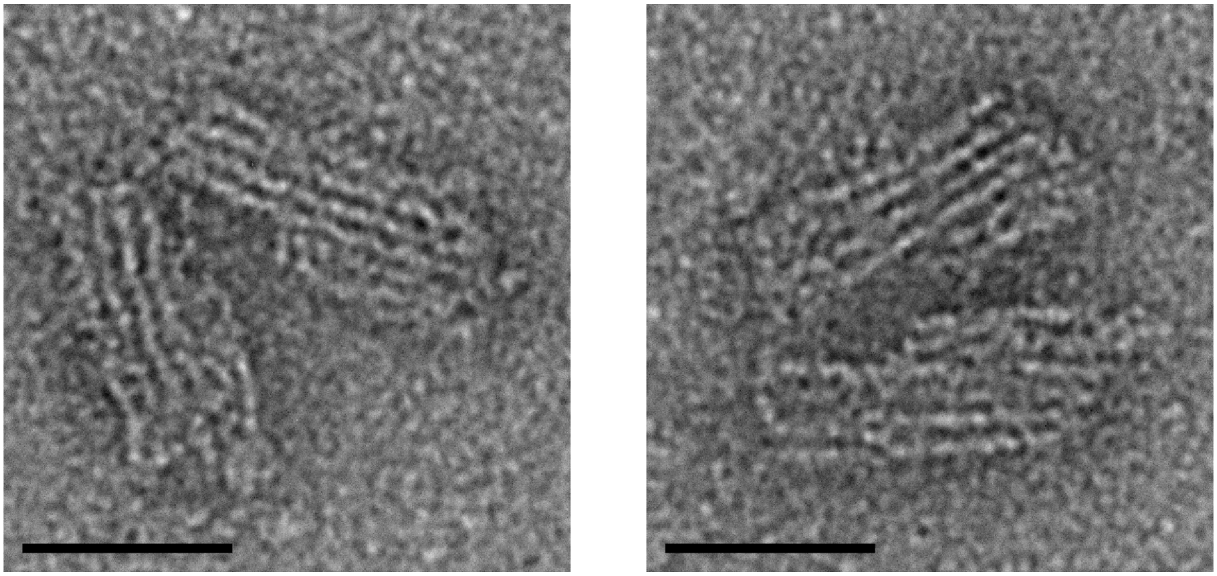

**Figure S2:** Exemplaric images of DNA origami structures after initial annealing according to Table S3 prior to incubation with the seam staples. Scale bars: 25 nm.

#### Dimer characterisation

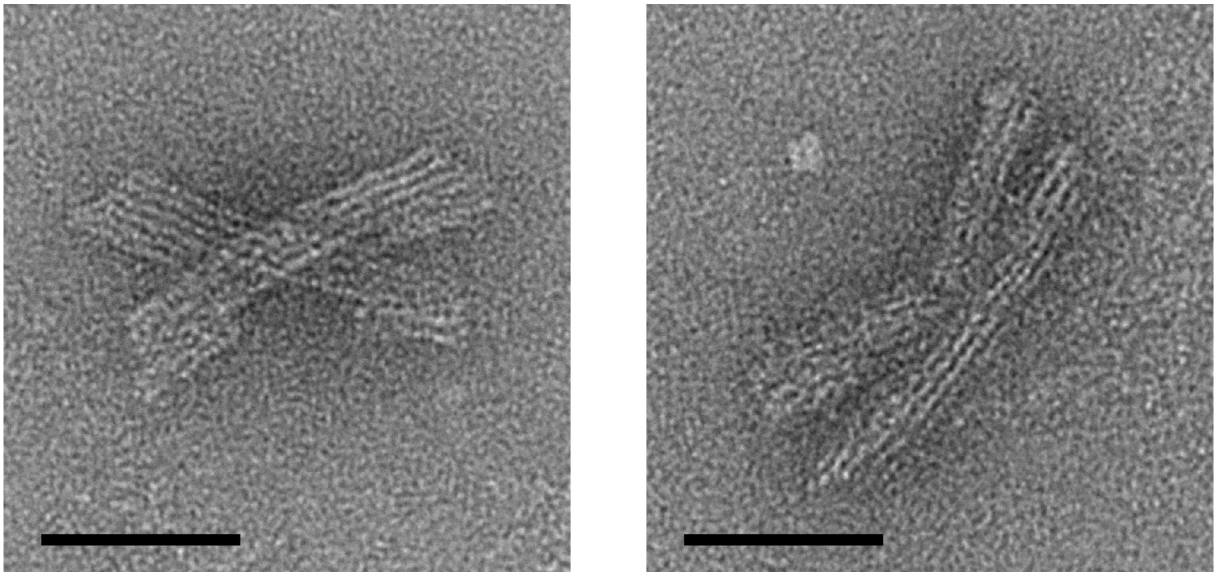

**Figure S3:** Exemplaric images of DNA origami structures extracted from the dimer band of a 38.8 pN structure mix. The structures showed either cross-like assemblies without apparent defects to the individual force clamps (left) or fusion structures with defects at the seam area (right). Scale bar: 50 nm.

##### DNA origamis structures after analysis

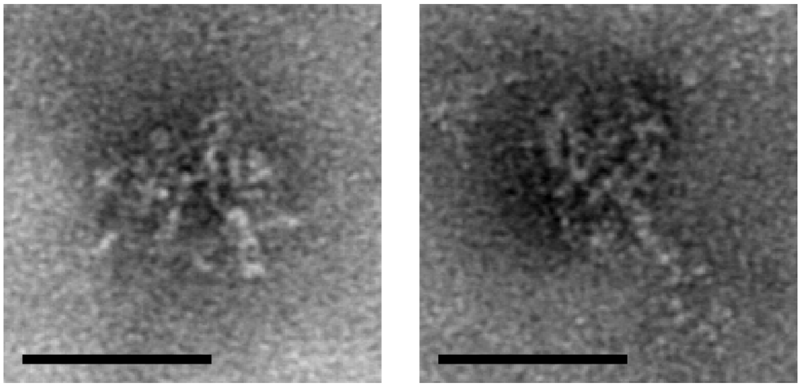

**Figure S4:** Exemplaric images of DNA origami force clamps extracted from a $Mg^{2+}$-free agarose after perfoming gel electrophoresis for 1.5 hours. Scale bars: 50 nm.

#### Enzymatic restriction of origamis

For every experiment, a batch solution comprising ultrapure $H_2O$ (Milli-Q®, Merck KGaA, Darmstadt; all further references of $H_2O$ refer to ultrapure $H_2O$), enzyme buffer concentrate and enzyme concentrate was prepared and aliquoted among the individually investigated forces. For EcoRV, a total of 8 replications were run; each replication included all forces and all times. Reactions were performed at 37℃. and were stopped with a quenching solution; immediately afterwards, the samples were transferred to 4℃. The samples were then subsequently transferred to a ice-cooled 1% agarose gel (Wide Mini-Sub Cell GT Cell, Bio-Rad Laboratories GmbH, Feldkirchen) using 1xTAE as running buffer. Gels ran at 70 V for 1.5 hours protected from light and were kept cool in an ice-water bath. Table S4 shows the total solution volumes; Table S5 describes the composition of the quenching solution. Adding the quenching solution resulted in an final EDTA concentration of 15 mM in the reaction mix, well above the total $Mg^{2+}$ concentration in the reaction mix of 12 mM, thus resulting in full chelation of $Mg^{2+}$ ions and subsequent termination of enzymatic activity.

| Component | Volume [µl] |
|---|---|
| $H_2O$ | 14 |
| enzyme buffer concentrate | 2 |
| enzyme concentrate (100 U/µl) | 2 |
| DNA origami solution (30 nM) | 2 |
| **Total (during reaction)** | **20** |
| Quenching solution | 4 |
| **Total (after reaction)** | **24** |

**Table S4:** Reaction mixture for every individual time point measurement

| Component | Volume [µl] |
|---|---|
| 6x loading buffer (Ficoll-based, with Orange G) | 760 |
| 0.5 M EDTA in $H_2O$ | 180 |
| 10 % SDS solution | 60 |
| **Total** | **1000** |

**Table S5:** Composition of quenching solution, scaled to 1 ml total volume.

Enzymes were ordered from New England Biolabs GmbH (Frankfurt, Germany). For EcoRV and BamHI, 100 U/µl equated to 1.0 µM; for EcoRI, 100 U/µl equated to 0.3 µM.

#### Data analysis

The agarose gels were subsequently imaged by using a GE Typhoon$^{TM}$ FLA 9000 gel imager. The band profiles were obtained using Fiji/ImageJ v1.53 by manually placing boxes of width 5.0 mm over each individual band and using the *Plot profile* tool to extract the average band intensity along the running direction of the agarose gel. Where necessary, images were rotated using the built-in ImageJ rotation tool using bicubic interpolation prior to band extraction to ensure proper alignment of boxes and bands. The fluorescent signal for each band was determined by subtracting the peak value from the average background. The average background was calculated by taking the average value of the data located 0.25 to 0.5 cm behind the peak (see Figure S5). For BamHI data, the Cy5 staple fragment was so close to the main band that its signal bleeded into the background range shown in Figure S5. We defined the background by taking the mean background of the four bands per replica that were not cut (i.e. the four t = 0s measurements for the four investigated forces). Based on the precision and homogeneity of laser gel readers, we do not expect any significant changes in the obtained results.

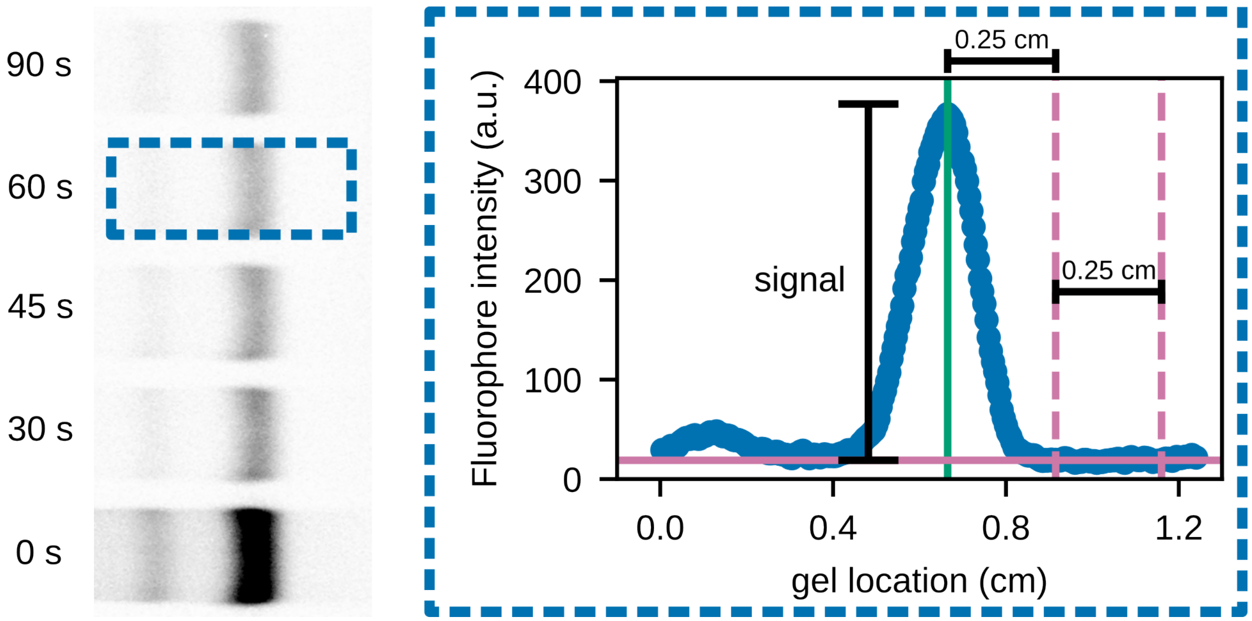


**Figure S5:** Data analysis from gel imaging. The Cy5 gel image of Figure 3c (left) is rotated by 90 degrees to place a signal extraction box (left blue box) for the *Plot profile* tool to take the average along the vertical axis, yielding yield an average band intensity profile (right blue box) along the horizontal travel direction during gel electrophoresis (left to right). The signal is then extracted from the difference between the maximum fluorescence intensity (located at the green line) and the background intensity (solid red line). The background intensity is determined by the average of the fluorescence intensity in a predefined region behind the maximum (dashed red lines).

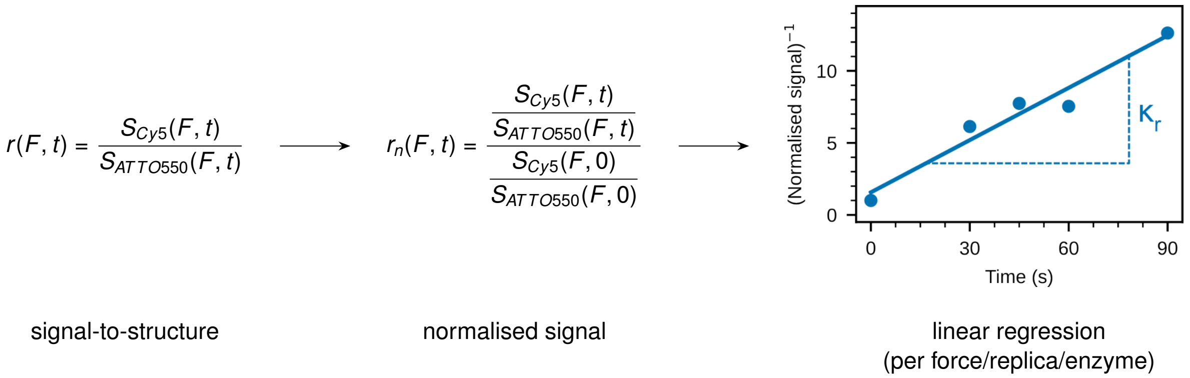


**Figure S6:** Visualisation of data pipeline from individual fluorescent signals to restriction rates.

To analyse the data (Figure S6), we calculate the signal-to-structure ratio $r$ by taking the Cy5 signal $S_{Cy5}$ of a specific force $F$ at a specific time $t$ as described in Figure S5 and dividing it by the ATTO550 signal $S_{ATTO550}$ of the same sample:

$$r(F, t) = \frac{S_{Cy5}(F, t)}{S_{ATTO550}(F, t)}$$

To normalise all force values with respect to the initial fluorophore signals (i.e. $t = 0s$), we divide the signal-to-structure ratios with the signal-to-structure ratio of the $t = 0s$ measurement:

$$r_n(F, t) = \frac{r_n(F, t)}{r_n(F, 0)}$$

We then plot $(r_n(F, t))^{-1}$ versus time for each individual force $F$ and use linear regression to determine the restriction rate $\kappa_r(F)$ for each force $F$, shown as the slope of each curve. For every enzyme, we obtained in total four (eight) restriction rates $\kappa_r(F)$; the statistical results of all $\kappa_r(F)$ values are shown in Figure 4.

#### BamHI restriction reaction data

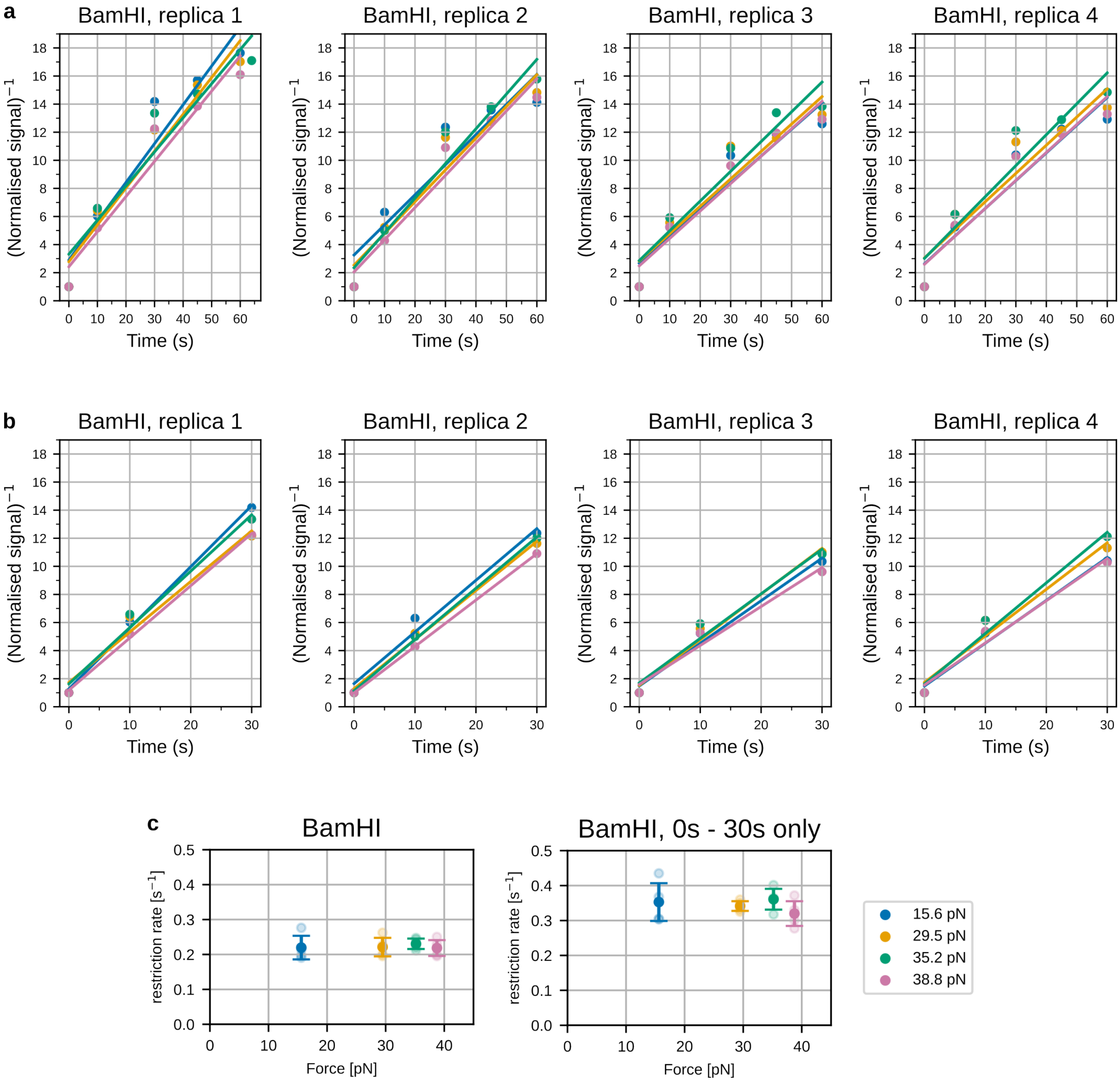


**Figure S7:** Enzymatic restriction of BamHI (legend for all plots in **c**). **a,** Replica data used to generate cumulative BamHI data in Figure 4. **b,** Analysis of data from **a** excluding all values with reaction times larger than 30 seconds. **c,** Comparison between datasets shown in **a** and **b**. The qualitative result of force-independent restriction activity for BamHI is consistently shown in both datasets.

#### Adapted force region

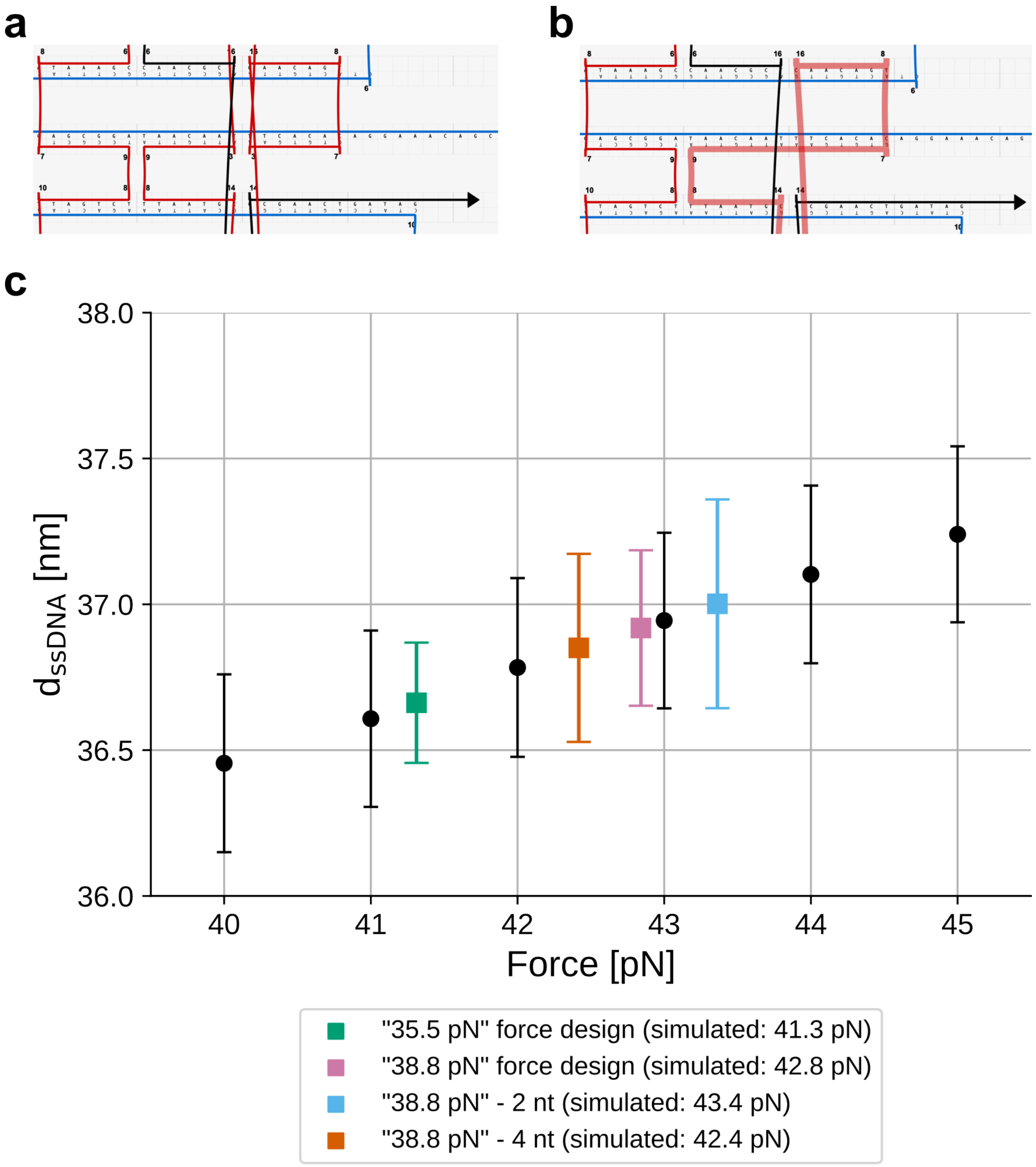


**Figure S8: Adapted staples design for force region. a,** Initial force staple design that was used for experiments in the paper. **b,** Adapted force staple design that increases the length of the staple domain exposed to the ssDNA region from 7 bp to 14 bp. **c,** Simulation data for structures with adapted force staple design including structures that have a shortened ssDNA region compared to the initial design (reduced by 2 (blue) and 4 (orange) nucleotides, respectively).

To increase rigidity in the inital staple layout of the structure (Figure S8a), two force staples have been adapted to expand the dsDNA domain that is directly exposed to the ssDNA strain from 7 nt to 14 nt (Figure S8b) to mitigate melting effects at the point of highest strain for the structure. When simulating the structures used for the experiments in Fig. 2-4 using the adapted force staple (Figure S8c), significant increases in force can be achieved. When shortening the ssDNA domain even further (i.e. further increasing the force applied by the entropic spring), we initially see a further increase in force up to 43 pN with slightly larger fluctuations in ssDNA length. We attribute this to force-induced melting of the force tower region adjacent to the entropic spring (bold red staple in **b**).

#### caDNAno design

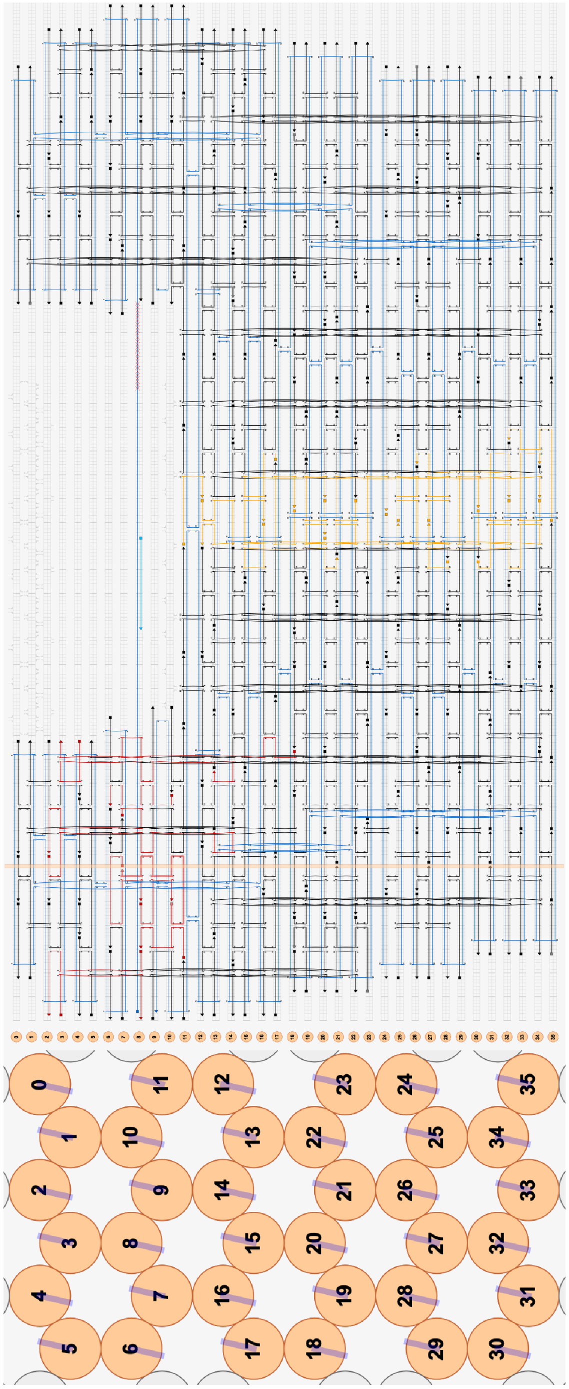

**Figure S9: caDNAno design of DNA origami force clamp.** The staples are split into three groups: core staples (black, constant for all forces), seam staples (yellow, constant for all forces) and force staples (red, individual sets for every force). All extruding staple ends have been modified to carry 4 T bases. The depicted structure shows the 15.6 pN structure with a 28 bp dsDNA region formed by the scaffold and the respective probing staples for every enzyme. To apply other forces, free nucleotides were removed to the left of the dsDNA strand region, shortening the total ssDNA region while keeping all non-red staples constant. The total number of free nucleotides for all force designs is shown in Table S1.

### DNA sequences

#### EcoRI/BamHI scaffold sequence (circular ssDNA, p8634 scaffold of M13mp18 bacteriophage)

CTCATTAGGCACCCCAGGCTTTACACTTTATGCTTCCGGCTCGTATGTTGTGTGGAATTGTGAGCGGATAA
CAATTTCACACAGGAAACAGCTATGACCATGATTACGAATTCGAGCTCGGTACCCGGGGATCCATTCTCCT
GTGACTCGGAAGTGCATTTATCATCTCCATAAAACAAAACCCGCCGTAGCGAGTTCAGATAAAATAAATCC
CCGCGAGTGCGAGGATTGTTATGTAATATTGGGTTTAATCATCTATATGTTTTGTACAGAGAGGGCAAGTAT
CGTTTCCACCGTACTCGTGATAATAATTTTGCACGGTATCAGTCATTTCTCGCACATTGCAGAATGGGGATT
TGTCTTCATTAGACTTATAAACCTTCATGGAATATTTGTATGCCGACTCTATATCTATACCTTCATCTACATA
AACACCTTCGTGATGTCTGCATGGAGACAAGACACCGGATCTGCACAACATTGATAACGCCCAATCTTTTT
GCTCAGACTCTAACTCATTGATACTCATTTATAAACTCCTTGCAATGTATGTCGTTTCAGCTAAACGGTATCA
GCAATGTTTATGTAAAGAAACAGTAAGATAATACTCAACCCGATGTTTGAGTACGGTCATCATCTGACACTA
CAGACTCTGGCATCGCTGTGAAGACGACGCGAAATTCAGCATTTTCACAAGCGTTATCTTTTACAAAACCG
ATCTCACTCTCCTTTGATGCGAATGCCAGCGTCAGACATCATATGCAGATACTCACCTGCATCCTGAACCC
ATTGACCTCCAACCCCGTAATAGCGATGCGTAATGATGTCGATAGTTACTAACGGGTCTTGTTCGATTAACT
GCCGCAGAAACTCTTCCAGGTCACCAGTGCAGTGCTTGATAACAGGAGTCTTCCCAGGATGGCGAACAAC
AAGAAACTGGTTTCCGTCTTCACGGACTTCGTTGCTTTCCAGTTTAGCAATACGCTTACTCCCATCCGAGAT
AACACCTTCGTAATACTCACGCTGCTCGTTGAGTTTTGATTTTGCTGTTTCAAGCTCAACACGCAGTTTCCC
TACTGTTAGCGCAATATCCTCGTTCTCCTGGTCGCGGCGTTTGATGTATTGCTGGTTTCTTTCCCGTTCATC
CAGCAGTTCCAGCACAATCGATGGTGTTACCAATTCATGGAAAAGGTCTGCGTCAAATCCCCAGTCGTCAT
GCATTGCCTGCTCTGCCGCTTCACGCAGTGCCTGAGAGTTAATTTCGCTCACTTCGAACCTCTCTGTTTAC
TGATAAGTTCCAGATCCTCCTGGCAACTTGCACAAGTCCGACAACCCTGAACGACCAGGCGTCTTCGTTCA
TCTATCGGATCGCCACACTCACAACAATGAGTGGCAGATATAGCCTGGTGGTTCAGGCGGCGCATTTTTAT
TGCTGTGTTGCGCTGTAATTCTTCTATTTCTGATGCTGAATCAATGATGTCTGCCATCTTTCATTAATCCCTG
AACTGTTGGTTAATACGCATGAGGGTGAATGCGAATAATAAAGCTTGGCACTGGCCGTCGTTTTACAACGT
CGTGACTGGGAAAACCCTGGCGTTACCCAACTTAATCGCCTTGCAGCACATCCCCCTTTCGCCAGCTGGC
GTAATAGCGAAGAGGCCCGCACCGATCGCCCTTCCCAACAGTTGCGCAGCCTGAATGGCGAATGGCGCT
TTGCCTGGTTTCCGGCACCAGAAGCGGTGCCGGAAAGCTGGCTGGAGTGCGATCTTCCTGAGGCCGATA
CTGTCGTCGTCCCCTCAAACTGGCAGATGCACGGTTACGATGCGCCCATCTACACCAACGTGACCTATCC
CATTACGGTCAATCCGCCGTTTGTTCCCACGGAGAATCCGACGGGTTGTTACTCGCTCACATTTAATGTTG
ATGAAAGCTGGCTACAGGAAGGCCAGACGCGAATTATTTTTGATGGCGTTCCTATTGGTTAAAAAATGAGC
TGATTTAACAAAAATTTAATGCGAATTTTAACAAAATATTAACGTTTACAATTTAAATATTTGCTTATACAATCT
TCCTGTTTTTGGGGCTTTTCTGATTATCAACCGGGGTACATATGATTGACATGCTAGTTTTACGATTACCGT
TCATCGATTCTCTTGTTTGCTCCAGACTCTCAGGCAATGACCTGATAGCCTTTGTAGATCTCTCAAAAATAG
CTACCCTCTCCGGCATTAATTTATCAGCTAGAACGGTTGAATATCATATTGATGGTGATTTGACTGTCTCCG
GCCTTTCTCACCCTTTTGAATCTTTACCTACACATTACTCAGGCATTGCATTTAAAATATATGAGGGTTCTAA
AAATTTTTATCCTTGCGTTGAAATAAAGGCTTCTCCCGCAAAAGTATTACAGGGTCATAATGTTTTTGGTACA
ACCGATTTAGCTTTATGCTCTGAGGCTTTATTGCTTAATTTTGCTAATTCTTTGCCTTGCCTGTATGATTTAT
TGGATGTTAATGCTACTACTATTAGTAGAATTGATGCCACCTTTTCAGCTCGCGCCCCAAATGAAAATATAG
CTAAACAGGTTATTGACCATTTGCGAAATGTATCTAATGGTCAAACTAAATCTACTCGTTCGCAGAATTGGG
AATCAACTGTTATATGGAATGAAACTTCCAGACACCGTACTTTAGTTGCATATTTAAAACATGTTGAGCTACA
GCATTATATTCAGCAATTAAGCTCTAAGCCATCCGCAAAAATGACCTCTTATCAAAAGGAGCAATTAAAGGT
ACTCTCTAATCCTGACCTGTTGGAGTTTGCTTCCGGTCTGGTTCGCTTTGAAGCTCGAATTAAAACGCGATA
TTTGAAGTCTTTCGGGCTTCCTCTTAATCTTTTTGATGCAATCCGCTTTGCTTCTGACTATAATAGTCAGGGT
AAAGACCTGATTTTTGATTTATGGTCATTCTCGTTTTCTGAACTGTTTAAAGCATTTGAGGGGGATTCAATGA
ATATTTATGACGATTCCGCAGTATTGGACGCTATCCAGTCTAAACATTTTACTATTACCCCCTCTGGCAAAA
CTTCTTTTGCAAAAGCCTCTCGCTATTTTGGTTTTTATCGTCGTCTGGTAAACGAGGGTTATGATAGTGTTG
CTCTTACTATGCCTCGTAATTCCTTTTGGCGTTATGTATCTGCATTAGTTGAATGTGGTATTCCTAAATCTCA
ACTGATGAATCTTTCTACCTGTAATAATGTTGTTCCGTTAGTTCGTTTTATTAACGTAGATTTTTCTTCCCAAC
GTCCTGACTGGTATAATGAGCCAGTTCTTAAAATCGCATAAGGTAATTCACAATGATTAAAGTTGAAATTAA
ACCATCTCAAGCCCAATTTACTACTCGTTCTGGTGTTTCTCGTCAGGGCAAGCCTTATTCACTGAATGAGCA
GCTTTGTTACGTTGATTTGGGTAATGAATATCCGGTTCTTGTCAAGATTACTCTTGATGAAGGTCAGCCAGC

CTATGCGCCTGGTCTGTACACCGTTCATCTGTCCTCTTTCAAAGTTGGTCAGTTCGGTTCCCTTATGATTGA
CCGTCTGCGCCTCGTTCCGGCTAAGTAACATGGAGCAGGTCGCGGATTTCGACACAATTTATCAGGCGAT
GATACAAATCTCCGTTGTACTTTGTTTCGCGCTTGGTATAATCGCTGGGGGTCAAAGATGAGTGTTTTAGTG
TATTCTTTTGCCTCTTTCGTTTTAGGTTGGTGCCTTCGTAGTGGCATTACGTATTTTACCCGTTTAATGGAAA
CTTCCTCATGAAAAAGTCTTTAGTCCTCAAAGCCTCTGTAGCCGTTGCTACCCTCGTTCCGATGCTGTCTTT
CGCTGCTGAGGGTGACGATCCCGCAAAAGCGGCCTTTAACTCCCTGCAAGCCTCAGCGACCGAATATATC
GGTTATGCGTGGGCGATGGTTGTTGTCATTGTCGGCGCAACTATCGGTATCAAGCTGTTTAAGAAATTCAC
CTCGAAAGCAAGCTGATAAACCGATACAATTAAAGGCTCCTTTTGGAGCCTTTTTTTTGGAGATTTTCAACG
TGAAAAAATTATTATTCGCAATTCCTTTAGTTGTTCCTTTCTATTCTCACTCCGCTGAAACTGTTGAAAGTTG
TTTAGCAAAATCCCATACAGAAAATTCATTTACTAACGTCTGGAAAGACGACAAAACTTTAGATCGTTACGC
TAACTATGAGGGCTGTCTGTGGAATGCTACAGGCGTTGTAGTTTGTACTGGTGACGAAACTCAGTGTTACG
GTACATGGGTTCCTATTGGGCTTGCTATCCCTGAAAATGAGGGTGGTGGCTCTGAGGGTGGCGGTTCTGA
GGGTGGCGGTTCTGAGGGTGGCGGTACTAAACCTCCTGAGTACGGTGATACACCTATTCCGGGCTATACT
TATATCAACCCTCTCGACGGCACTTATCCGCCTGGTACTGAGCAAAACCCCGCTAATCCTAATCCTTCTCTT
GAGGAGTCTCAGCCTCTTAATACTTTCATGTTTCAGAATAATAGGTTCCGAAATAGGCAGGGGGCATTAAC
TGTTTATACGGGCACTGTTACTCAAGGCACTGACCCCGTTAAAACTTATTACCAGTACACTCCTGTATCATC
AAAAGCCATGTATGACGCTTACTGGAACGGTAAATTCAGAGACTGCGCTTTCCATTCTGGCTTTAATGAGG
ATTTATTTGTTTGTGAATATCAAGGCCAATCGTCTGACCTGCCTCAACCTCCTGTCAATGCTGGCGGCGGC
TCTGGTGGTGGTTCTGGTGGCGGCTCTGAGGGTGGTGGCTCTGAGGGTGGCGGTTCTGAGGGTGGCGG
CTCTGAGGGAGGCGGTTCCGGTGGTGGCTCTGGTTCCGGTGATTTTGATTATGAAAAGATGGCAAACGCT
AATAAGGGGGCTATGACCGAAAATGCCGATGAAAACGCGCTACAGTCTGACGCTAAAGGCAAACTTGATT
CTGTCGCTACTGATTACGGTGCTGCTATCGATGGTTTCATTGGTGACGTTTCCGGCCTTGCTAATGGTAAT
GGTGCTACTGGTGATTTTGCTGGCTCTAATTCCCAAATGGCTCAAGTCGGTGACGGTGATAATTCACCTTT
AATGAATAATTTCCGTCAATATTTACCTTCCCTCCCTCAATCGGTTGAATGTCGCCCTTTTGTCTTTGGCGC
TGGTAAACCATATGAATTTTCTATTGATTGTGACAAAATAAACTTATTCCGTGGTGTCTTTGCGTTTCTTTTAT
ATGTTGCCACCTTTATGTATGTATTTTCTACGTTTGCTAACATACTGCGTAATAAGGAGTCTTAATCATGCCA
GTTCTTTTGGGTATTCCGTTATTATTGCGTTTCCTCGGTTTCCTTCTGGTAACTTTGTTCGGCTATCTGCTTA
CTTTTCTTAAAAAGGGCTTCGGTAAGATAGCTATTGCTATTTCATTGTTTCTTGCTCTTATTATTGGGCTTAA
CTCAATTCTTGTGGGTTATCTCTCTGATATTAGCGCTCAATTACCCTCTGACTTTGTTCAGGGTGTTCAGTTA
ATTCTCCCGTCTAATGCGCTTCCCTGTTTTTATGTTATTCTCTCTGTAAAGGCTGCTATTTTCATTTTTGACG
TTAAACAAAAAATCGTTTCTTATTTGGATTGGGATAAATAATATGGCTGTTTATTTTGTAACTGGCAAATTAG
GCTCTGGAAAGACGCTCGTTAGCGTTGGTAAGATTCAGGATAAAATTGTAGCTGGGTGCAAAATAGCAACT
AATCTTGATTTAAGGCTTCAAAACCTCCCGCAAGTCGGGAGGTTCGCTAAAACGCCTCGCGTTCTTAGAAT
ACCGGATAAGCCTTCTATATCTGATTTGCTTGCTATTGGGCGCGGTAATGATTCCTACGATGAAAATAAAAA
CGGCTTGCTTGTTCTCGATGAGTGCGGTACTTGGTTTAATACCCGTTCTTGGAATGATAAGGAAAGACAGC
CGATTATTGATTGGTTTCTACATGCTCGTAAATTAGGATGGGATATTATTTTTCTTGTTCAGGACTTATCTAT
TGTTGATAAACAGGCGCGTTCTGCATTAGCTGAACATGTTGTTTATTGTCGTCGTCTGGACAGAATTACTTT
ACCTTTTGTCGGTACTTTATATTCTCTTATTACTGGCTCGAAAATGCCTCTGCCTAAATTACATGTTGGCGTT
GTTAAATATGGCGATTCTCAATTAAGCCCTACTGTTGAGCGTTGGCTTTATACTGGTAAGAATTTGTATAAC
GCATATGATACTAAACAGGCTTTTTCTAGTAATTATGATTCCGGTGTTTATTCTTATTTAACGCCTTATTTATC
ACACGGTCGGTATTTCAAACCATTAAATTTAGGTCAGAAGATGAAATTAACTAAAATATATTTGAAAAAGTTT
TCTCGCGTTCTTTGTCTTGCGATTGGATTTGCATCAGCATTTACATATAGTTATATAACCCAACCTAAGCCG
GAGGTTAAAAAGGTAGTCTCTCAGACCTATGATTTTGATAAATTCACTATTGACTCTTCTCAGCGTCTTAATC
TAAGCTATCGCTATGTTTTCAAGGATTCTAAGGGAAAATTAATTAATAGCGACGATTTACAGAAGCAAGGTT
ATTCACTCACATATATTGATTTATGTACTGTTTCCATTAAAAAAGGTAATTCAAATGAAATTGTTAAATGTAAT
TAATTTTGTTTTCTTGATGTTTGTTTCATCATCTTCTTTTGCTCAGGTAATTGAAATGAATAATTCGCCTCTGC
GCGATTTTGTAACTTGGTATTCAAAGCAATCAGGCGAATCCGTTATTGTTTCTCCCGATGTAAAAGGTACTG
TTACTGTATATTCATCTGACGTTAAACCTGAAAATCTACGCAATTTCTTTATTTCTGTTTTACGTGCAAATAAT
TTTGATATGGTAGGTTCTAACCCTTCCATTATTCAGAAGTATAATCCAAACAATCAGGATTATATTGATGAAT
TGCCATCATCTGATAATCAGGAATATGATGATAATTCCGCTCCTTCTGGTGGTTTCTTTGTTCCGCAAAATG
ATAATGTTACTCAAACTTTTAAAATTAATAACGTTCGGGCAAAGGATTTAATACGAGTTGTCGAATTGTTTGT
AAAGTCTAATACTTCTAAATCCTCAAATGTATTATCTATTGACGGCTCTAATCTATTAGTTGTTAGTGCTCCT
AAAGATATTTTAGATAACCTTCCTCAATTCCTTTCAACTGTTGATTTGCCAACTGACCAGATATTGATTGAGG
GTTTGATATTTGAGGTTCAGCAAGGTGATGCTTTAGATTTTTCATTTGCTGCTGGCTCTCAGCGTGGCACTG

TTGCAGGCGGTGTTAATACTGACCGCCTCACCTCTGTTTTATCTTCTGCTGGTGGTTCGTTCGGTATTTTTA
ATGGCGATGTTTTAGGGCTATCAGTTCGCGCATTAAAGACTAATAGCCATTCAAAAATATTGTCTGTGCCAC
GTATTCTTACGCTTTCAGGTCAGAAGGGTTCTATCTCTGTTGGCCAGAATGTCCCTTTTATTACTGGTCGTG
TGACTGGTGAATCTGCCAATGTAAATAATCCATTTCAGACGATTGAGCGTCAAAATGTAGGTATTTCCATGA
GCGTTTTTCCTGTTGCAATGGCTGGCGGTAATATTGTTCTGGATATTACCAGCAAGGCCGATAGTTTGAGT
TCTTCTACTCAGGCAAGTGATGTTATTACTAATCAAAGAAGTATTGCTACAACGGTTAATTTGCGTGATGGA
CAGACTCTTTTACTCGGTGGCCTCACTGATTATAAAAACACTTCTCAGGATTCTGGCGTACCGTTCCTGTCT
AAAATCCCTTTAATCGGCCTCCTGTTTAGCTCCCGCTCTGATTCTAACGAGGAAAGCACGTTATACGTGCTC
GTCAAAGCAACCATAGTACGCGCCCTGTAGCGGCGCATTAAGCGCGGCGGGTGTGGTGGTTACGCGCAG
CGTGACCGCTACACTTGCCAGCGCCCTAGCGCCCGCTCCTTTCGCTTTCTTCCCTTCCTTTCTCGCCACGT
TCGCCGGCTTTCCCCGTCAAGCTCTAAATCGGGGGCTCCCTTTAGGGTTCCGATTTAGTGCTTTACGGCAC
CTCGACCCCAAAAAACTTGATTTGGGTGATGGTTCACGTAGTGGGCCATCGCCCTGATAGACGGTTTTTCG
CCCTTTGACGTTGGAGTCCACGTTCTTTAATAGTGGACTCTTGTTCCAAACTGGAACAACACTCAACCCTAT
CTCGGGCTATTCTTTTGATTTATAAGGGATTTTGCCGATTTCGGAACCACCATCAAACAGGATTTTCGCCTG
CTGGGGCAAACCAGCGTGGACCGCTTGCTGCAACTCTCTCAGGGCCAGGCGGTGAAGGGCAATCAGCTG
TTGCCCGTCTCACTGGTGAAAAGAAAAACCACCCTGGCGCCCAATACGCAAACCGCCTCTCCCCGCGCGT
TGGCCGATTCATTAATGCAGCTGGCACGACAGGTTTCCCGACTGGAAAGCGGGCAGTGAGCGCAACGCA
ATTAATGTGAGTTAGCTCA

#### EcoRV scaffold preparation

The sequence was created by double-digesting the EcoRI/BamHI scaffold using EcoRI and BamHI and cloning in a sequence containing the EcoRV restriction site GATATC using a standard cloning protocol as described earlier [1]. Correct cloning was verified via sequencing (Mix2Seq, Eurofins Genomics GmbH, Ebersberg, Germany).

Cut out sequence (between BamHI and EcoRI):

```
  scaffold:    5’ - AATT CGAGCT CGGTACC CGGG      - 3’
complement:    3’ -      GCTCGA GCCATGG GCCC CTAG - 5’
```

Replacement sequence with EcoRV restriction site (marked in red):

```
  scaffold:    5’ - AATT GATATC TCCTCCT CCTC      - 3’
complement:    3’ -      CTATAG AGGAGGA GGAG CTAG - 5’
```

##### Full sequence after cloning

CTCATTAGGCACCCCAGGCTTTACACTTTATGCTTCCGGCTCGTATGTTGTGTGGAATTGTGAGCGGATAA
CAATTTCACACAGGAAACAGCTATGACCATGATTACGAATTGATATCTCCTCCTCCTCGATCCATTCTCCTG
TGACTCGGAAGTGCATTTATCATCTCCATAAAACAAAACCCGCCGTAGCGAGTTCAGATAAAATAAATCCCC
GCGAGTGCGAGGATTGTTATGTAATATTGGGTTTAATCATCTATATGTTTTGTACAGAGAGGGCAAGTATCG
TTTCCACCGTACTCGTGATAATAATTTTGCACGGTATCAGTCATTTCTCGCACATTGCAGAATGGGGATTTG
TCTTCATTAGACTTATAAACCTTCATGGAATATTTGTATGCCGACTCTATATCTATACCTTCATCTACATAAAC
ACCTTCGTGATGTCTGCATGGAGACAAGACACCGGATCTGCACAACATTGATAACGCCCAATCTTTTTGCT
CAGACTCTAACTCATTGATACTCATTTATAAACTCCTTGCAATGTATGTCGTTTCAGCTAAACGGTATCAGCA
ATGTTTATGTAAAGAAACAGTAAGATAATACTCAACCCGATGTTTGAGTACGGTCATCATCTGACACTACAG
ACTCTGGCATCGCTGTGAAGACGACGCGAAATTCAGCATTTTCACAAGCGTTATCTTTTACAAAACCGATCT
CACTCTCCTTTGATGCGAATGCCAGCGTCAGACATCATATGCAGATACTCACCTGCATCCTGAACCCATTG
ACCTCCAACCCCGTAATAGCGATGCGTAATGATGTCGATAGTTACTAACGGGTCTTGTTCGATTAACTGCC
GCAGAAACTCTTCCAGGTCACCAGTGCAGTGCTTGATAACAGGAGTCTTCCCAGGATGGCGAACAACAAG
AAACTGGTTTCCGTCTTCACGGACTTCGTTGCTTTCCAGTTTAGCAATACGCTTACTCCCATCCGAGATAAC
ACCTTCGTAATACTCACGCTGCTCGTTGAGTTTTGATTTTGCTGTTTCAAGCTCAACACGCAGTTTCCCTAC
TGTTAGCGCAATATCCTCGTTCTCCTGGTCGCGGCGTTTGATGTATTGCTGGTTTCTTTCCCGTTCATCCAG
CAGTTCCAGCACAATCGATGGTGTTACCAATTCATGGAAAAGGTCTGCGTCAAATCCCCAGTCGTCATGCA
TTGCCTGCTCTGCCGCTTCACGCAGTGCCTGAGAGTTAATTTCGCTCACTTCGAACCTCTCTGTTTACTGAT
AAGTTCCAGATCCTCCTGGCAACTTGCACAAGTCCGACAACCCTGAACGACCAGGCGTCTTCGTTCATCTA

TCGGATCGCCACACTCACAACAATGAGTGGCAGATATAGCCTGGTGGTTCAGGCGGCGCATTTTTATTGCT
GTGTTGCGCTGTAATTCTTCTATTTCTGATGCTGAATCAATGATGTCTGCCATCTTTCATTAATCCCTGAACT
GTTGGTTAATACGCATGAGGGTGAATGCGAATAATAAAGCTTGGCACTGGCCGTCGTTTTACAACGTCGTG
ACTGGGAAAACCCTGGCGTTACCCAACTTAATCGCCTTGCAGCACATCCCCCTTTCGCCAGCTGGCGTAAT
AGCGAAGAGGCCCGCACCGATCGCCCTTCCCAACAGTTGCGCAGCCTGAATGGCGAATGGCGCTTTGCC
TGGTTTCCGGCACCAGAAGCGGTGCCGGAAAGCTGGCTGGAGTGCGATCTTCCTGAGGCCGATACTGTC
GTCGTCCCCTCAAACTGGCAGATGCACGGTTACGATGCGCCCATCTACACCAACGTGACCTATCCCATTAC
GGTCAATCCGCCGTTTGTTCCCACGGAGAATCCGACGGGTTGTTACTCGCTCACATTTAATGTTGATGAAA
GCTGGCTACAGGAAGGCCAGACGCGAATTATTTTTGATGGCGTTCCTATTGGTTAAAAAATGAGCTGATTT
AACAAAAATTTAATGCGAATTTTAACAAAATATTAACGTTTACAATTTAAATATTTGCTTATACAATCTTCCTG
TTTTTGGGGCTTTTCTGATTATCAACCGGGGTACATATGATTGACATGCTAGTTTTACGATTACCGTTCATC
GATTCTCTTGTTTGCTCCAGACTCTCAGGCAATGACCTGATAGCCTTTGTAGATCTCTCAAAAATAGCTACC
CTCTCCGGCATTAATTTATCAGCTAGAACGGTTGAATATCATATTGATGGTGATTTGACTGTCTCCGGCCTT
TCTCACCCTTTTGAATCTTTACCTACACATTACTCAGGCATTGCATTTAAAATATATGAGGGTTCTAAAAATT
TTTATCCTTGCGTTGAAATAAAGGCTTCTCCCGCAAAAGTATTACAGGGTCATAATGTTTTTGGTACAACCG
ATTTAGCTTTATGCTCTGAGGCTTTATTGCTTAATTTTGCTAATTCTTTGCCTTGCCTGTATGATTTATTGGAT
GTTAATGCTACTACTATTAGTAGAATTGATGCCACCTTTTCAGCTCGCGCCCCAAATGAAAATATAGCTAAA
CAGGTTATTGACCATTTGCGAAATGTATCTAATGGTCAAACTAAATCTACTCGTTCGCAGAATTGGGAATCA
ACTGTTATATGGAATGAAACTTCCAGACACCGTACTTTAGTTGCATATTTAAAACATGTTGAGCTACAGCATT
ATATTCAGCAATTAAGCTCTAAGCCATCCGCAAAAATGACCTCTTATCAAAAGGAGCAATTAAAGGTACTCT
CTAATCCTGACCTGTTGGAGTTTGCTTCCGGTCTGGTTCGCTTTGAAGCTCGAATTAAAACGCGATATTTGA
AGTCTTTCGGGCTTCCTCTTAATCTTTTTGATGCAATCCGCTTTGCTTCTGACTATAATAGTCAGGGTAAAG
ACCTGATTTTTGATTTATGGTCATTCTCGTTTTCTGAACTGTTTAAAGCATTTGAGGGGGATTCAATGAATAT
TTATGACGATTCCGCAGTATTGGACGCTATCCAGTCTAAACATTTTACTATTACCCCCTCTGGCAAAACTTC
TTTTGCAAAAGCCTCTCGCTATTTTGGTTTTTATCGTCGTCTGGTAAACGAGGGTTATGATAGTGTTGCTCT
TACTATGCCTCGTAATTCCTTTTGGCGTTATGTATCTGCATTAGTTGAATGTGGTATTCCTAAATCTCAACTG
ATGAATCTTTCTACCTGTAATAATGTTGTTCCGTTAGTTCGTTTTATTAACGTAGATTTTTCTTCCCAACGTCC
TGACTGGTATAATGAGCCAGTTCTTAAAATCGCATAAGGTAATTCACAATGATTAAAGTTGAAATTAAACCA
TCTCAAGCCCAATTTACTACTCGTTCTGGTGTTTCTCGTCAGGGCAAGCCTTATTCACTGAATGAGCAGCTT
TGTTACGTTGATTTGGGTAATGAATATCCGGTTCTTGTCAAGATTACTCTTGATGAAGGTCAGCCAGCCTAT
GCGCCTGGTCTGTACACCGTTCATCTGTCCTCTTTCAAAGTTGGTCAGTTCGGTTCCCTTATGATTGACCG
TCTGCGCCTCGTTCCGGCTAAGTAACATGGAGCAGGTCGCGGATTTCGACACAATTTATCAGGCGATGATA
CAAATCTCCGTTGTACTTTGTTTCGCGCTTGGTATAATCGCTGGGGGTCAAAGATGAGTGTTTTAGTGTATT
CTTTTGCCTCTTTCGTTTTAGGTTGGTGCCTTCGTAGTGGCATTACGTATTTTACCCGTTTAATGGAAACTTC
CTCATGAAAAAGTCTTTAGTCCTCAAAGCCTCTGTAGCCGTTGCTACCCTCGTTCCGATGCTGTCTTTCGCT
GCTGAGGGTGACGATCCCGCAAAAGCGGCCTTTAACTCCCTGCAAGCCTCAGCGACCGAATATATCGGTT
ATGCGTGGGCGATGGTTGTTGTCATTGTCGGCGCAACTATCGGTATCAAGCTGTTTAAGAAATTCACCTCG
AAAGCAAGCTGATAAACCGATACAATTAAAGGCTCCTTTTGGAGCCTTTTTTTTGGAGATTTTCAACGTGAA
AAAATTATTATTCGCAATTCCTTTAGTTGTTCCTTTCTATTCTCACTCCGCTGAAACTGTTGAAAGTTGTTTAG
CAAAATCCCATACAGAAAATTCATTTACTAACGTCTGGAAAGACGACAAAACTTTAGATCGTTACGCTAACT
ATGAGGGCTGTCTGTGGAATGCTACAGGCGTTGTAGTTTGTACTGGTGACGAAACTCAGTGTTACGGTACA
TGGGTTCCTATTGGGCTTGCTATCCCTGAAAATGAGGGTGGTGGCTCTGAGGGTGGCGGTTCTGAGGGTG
GCGGTTCTGAGGGTGGCGGTACTAAACCTCCTGAGTACGGTGATACACCTATTCCGGGCTATACTTATATC
AACCCTCTCGACGGCACTTATCCGCCTGGTACTGAGCAAAACCCCGCTAATCCTAATCCTTCTCTTGAGGA
GTCTCAGCCTCTTAATACTTTCATGTTTCAGAATAATAGGTTCCGAAATAGGCAGGGGGCATTAACTGTTTA
TACGGGCACTGTTACTCAAGGCACTGACCCCGTTAAAACTTATTACCAGTACACTCCTGTATCATCAAAAGC
CATGTATGACGCTTACTGGAACGGTAAATTCAGAGACTGCGCTTTCCATTCTGGCTTTAATGAGGATTTATT
TGTTTGTGAATATCAAGGCCAATCGTCTGACCTGCCTCAACCTCCTGTCAATGCTGGCGGCGGCTCTGGTG
GTGGTTCTGGTGGCGGCTCTGAGGGTGGTGGCTCTGAGGGTGGCGGTTCTGAGGGTGGCGGCTCTGAG
GGAGGCGGTTCCGGTGGTGGCTCTGGTTCCGGTGATTTTGATTATGAAAAGATGGCAAACGCTAATAAGG
GGGCTATGACCGAAAATGCCGATGAAAACGCGCTACAGTCTGACGCTAAAGGCAAACTTGATTCTGTCGC
TACTGATTACGGTGCTGCTATCGATGGTTTCATTGGTGACGTTTCCGGCCTTGCTAATGGTAATGGTGCTA
CTGGTGATTTTGCTGGCTCTAATTCCCAAATGGCTCAAGTCGGTGACGGTGATAATTCACCTTTAATGAATA
ATTTCCGTCAATATTTACCTTCCCTCCCTCAATCGGTTGAATGTCGCCCTTTTGTCTTTGGCGCTGGTAAAC

CATATGAATTTTCTATTGATTGTGACAAAATAAACTTATTCCGTGGTGTCTTTGCGTTTCTTTTATATGTTGCC
ACCTTTATGTATGTATTTTCTACGTTTGCTAACATACTGCGTAATAAGGAGTCTTAATCATGCCAGTTCTTTT
GGGTATTCCGTTATTATTGCGTTTCCTCGGTTTCCTTCTGGTAACTTTGTTCGGCTATCTGCTTACTTTTCTT
AAAAAGGGCTTCGGTAAGATAGCTATTGCTATTTCATTGTTTCTTGCTCTTATTATTGGGCTTAACTCAATTC
TTGTGGGTTATCTCTCTGATATTAGCGCTCAATTACCCTCTGACTTTGTTCAGGGTGTTCAGTTAATTCTCC
CGTCTAATGCGCTTCCCTGTTTTTATGTTATTCTCTCTGTAAAGGCTGCTATTTTCATTTTTGACGTTAAACA
AAAAATCGTTTCTTATTTGGATTGGGATAAATAATATGGCTGTTTATTTTGTAACTGGCAAATTAGGCTCTGG
AAAGACGCTCGTTAGCGTTGGTAAGATTCAGGATAAAATTGTAGCTGGGTGCAAAATAGCAACTAATCTTG
ATTTAAGGCTTCAAAACCTCCCGCAAGTCGGGAGGTTCGCTAAAACGCCTCGCGTTCTTAGAATACCGGAT
AAGCCTTCTATATCTGATTTGCTTGCTATTGGGCGCGGTAATGATTCCTACGATGAAAATAAAAACGGCTTG
CTTGTTCTCGATGAGTGCGGTACTTGGTTTAATACCCGTTCTTGGAATGATAAGGAAAGACAGCCGATTATT
GATTGGTTTCTACATGCTCGTAAATTAGGATGGGATATTATTTTTCTTGTTCAGGACTTATCTATTGTTGATA
AACAGGCGCGTTCTGCATTAGCTGAACATGTTGTTTATTGTCGTCGTCTGGACAGAATTACTTTACCTTTTG
TCGGTACTTTATATTCTCTTATTACTGGCTCGAAAATGCCTCTGCCTAAATTACATGTTGGCGTTGTTAAATA
TGGCGATTCTCAATTAAGCCCTACTGTTGAGCGTTGGCTTTATACTGGTAAGAATTTGTATAACGCATATGA
TACTAAACAGGCTTTTTCTAGTAATTATGATTCCGGTGTTTATTCTTATTTAACGCCTTATTTATCACACGGTC
GGTATTTCAAACCATTAAATTTAGGTCAGAAGATGAAATTAACTAAAATATATTTGAAAAAGTTTTCTCGCGT
TCTTTGTCTTGCGATTGGATTTGCATCAGCATTTACATATAGTTATATAACCCAACCTAAGCCGGAGGTTAA
AAAGGTAGTCTCTCAGACCTATGATTTTGATAAATTCACTATTGACTCTTCTCAGCGTCTTAATCTAAGCTAT
CGCTATGTTTTCAAGGATTCTAAGGGAAAATTAATTAATAGCGACGATTTACAGAAGCAAGGTTATTCACTC
ACATATATTGATTTATGTACTGTTTCCATTAAAAAAGGTAATTCAAATGAAATTGTTAAATGTAATTAATTTTG
TTTTCTTGATGTTTGTTTCATCATCTTCTTTTGCTCAGGTAATTGAAATGAATAATTCGCCTCTGCGCGATTTT
GTAACTTGGTATTCAAAGCAATCAGGCGAATCCGTTATTGTTTCTCCCGATGTAAAAGGTACTGTTACTGTA
TATTCATCTGACGTTAAACCTGAAAATCTACGCAATTTCTTTATTTCTGTTTTACGTGCAAATAATTTTGATAT
GGTAGGTTCTAACCCTTCCATTATTCAGAAGTATAATCCAAACAATCAGGATTATATTGATGAATTGCCATCA
TCTGATAATCAGGAATATGATGATAATTCCGCTCCTTCTGGTGGTTTCTTTGTTCCGCAAAATGATAATGTTA
CTCAAACTTTTAAAATTAATAACGTTCGGGCAAAGGATTTAATACGAGTTGTCGAATTGTTTGTAAAGTCTAA
TACTTCTAAATCCTCAAATGTATTATCTATTGACGGCTCTAATCTATTAGTTGTTAGTGCTCCTAAAGATATTT
TAGATAACCTTCCTCAATTCCTTTCAACTGTTGATTTGCCAACTGACCAGATATTGATTGAGGGTTTGATATT
TGAGGTTCAGCAAGGTGATGCTTTAGATTTTTCATTTGCTGCTGGCTCTCAGCGTGGCACTGTTGCAGGCG
GTGTTAATACTGACCGCCTCACCTCTGTTTTATCTTCTGCTGGTGGTTCGTTCGGTATTTTTAATGGCGATG
TTTTAGGGCTATCAGTTCGCGCATTAAAGACTAATAGCCATTCAAAAATATTGTCTGTGCCACGTATTCTTAC
GCTTTCAGGTCAGAAGGGTTCTATCTCTGTTGGCCAGAATGTCCCTTTTATTACTGGTCGTGTGACTGGTG
AATCTGCCAATGTAAATAATCCATTTCAGACGATTGAGCGTCAAAATGTAGGTATTTCCATGAGCGTTTTTC
CTGTTGCAATGGCTGGCGGTAATATTGTTCTGGATATTACCAGCAAGGCCGATAGTTTGAGTTCTTCTACTC
AGGCAAGTGATGTTATTACTAATCAAAGAAGTATTGCTACAACGGTTAATTTGCGTGATGGACAGACTCTTT
TACTCGGTGGCCTCACTGATTATAAAAACACTTCTCAGGATTCTGGCGTACCGTTCCTGTCTAAAATCCCTT
TAATCGGCCTCCTGTTTAGCTCCCGCTCTGATTCTAACGAGGAAAGCACGTTATACGTGCTCGTCAAAGCA
ACCATAGTACGCGCCCTGTAGCGGCGCATTAAGCGCGGCGGGTGTGGTGGTTACGCGCAGCGTGACCGC
TACACTTGCCAGCGCCCTAGCGCCCGCTCCTTTCGCTTTCTTCCCTTCCTTTCTCGCCACGTTCGCCGGCT
TTCCCCGTCAAGCTCTAAATCGGGGGCTCCCTTTAGGGTTCCGATTTAGTGCTTTACGGCACCTCGACCCC
AAAAAACTTGATTTGGGTGATGGTTCACGTAGTGGGCCATCGCCCTGATAGACGGTTTTTCGCCCTTTGAC
GTTGGAGTCCACGTTCTTTAATAGTGGACTCTTGTTCCAAACTGGAACAACACTCAACCCTATCTCGGGCTA
TTCTTTTGATTTATAAGGGATTTTGCCGATTTCGGAACCACCATCAAACAGGATTTTCGCCTGCTGGGGCAA
ACCAGCGTGGACCGCTTGCTGCAACTCTCTCAGGGCCAGGCGGTGAAGGGCAATCAGCTGTTGCCCGTC
TCACTGGTGAAAAGAAAAACCACCCTGGCGCCCAATACGCAAACCGCCTCTCCCCGCGCGTTGGCCGATT
CATTAATGCAGCTGGCACGACAGGTTTCCCGACTGGAAAGCGGGCAGTGAGCGCAACGCAATTAATGTGA
GTTAGCTCA

#### Core staples

Every line represents the sequence of one staple.

TAATCGCGCTGCTCATTCAGTTGATAAAAAAAGAACATAACC

TGTCATCACGAAGAGGAAGCCCTGGAGAATGTGTTTAG
TAAGAGCAAGAAACTGTAGCGGAGCGGGATTAAGACCTT
TTTTAGATTTAGTCGTGAAGTGGAAAGAAGACTCCTCGCCGTCAAAGTGAAC
GGAGTAAGAGAACGAGGATATACCGCCA
TTTCAGGCAAAAACAGGAAGATGATAATGAACGGTTGCAGGTCATT
GCGAGAAACGAGAATTAACTGCCCACAA
CGCGGGGTTACTTGCCCTCTCTGTAGCT
TGCTACATTGCTGATACGTCAGATGATGACCGTAAGAGCGAT
ACTCAACATTAAATCTGAGAGGGGTAGCCTAG
TATTACGCCATAATGTGTATAGTTAGCGTAAAATAGAAAGGC
TAACGACATTGAATAAGTTTATTTAAGGTGGCATTACCCATCGATACCA
CCTAACAGTTTGCCTTGAGTAACACGTCATATCTCTGAGGTG
TAAACAGTACAGGGCGCGTCATGGCAAGAATGAAAAGTTACCTCTCCGCCCCA
AAAAGTAAAAACCATAAATCAAAAAAAGATTAAGG
AGTTGGCACCTTGCTGAACCTAACACCG
CGAGAGCGGGGACAGGATACCAGTAATAAAA
ATATGTCACACAAAGACAAAAGGGATATTGAGTCA
AGCATGGCTACGAACGAGTTTATCTATCCCATCCTAAT
TCAGGGACCGCCACAGGCTTGAGCATCGAGGACTAAAGACTTGGCA
ATAATCAACGCAAATAGAAGAACTCAAAAAA
GTCCATCGTGAGGCTCTGACCTGAAATATTACTAGGGAATTA
GGTAAATTTTAGCCTTTTTATCTGATTGTTGAGGATCTGGCGCAA
AAGTGTTATTTTATAAACGATTTTTTGTAGGGAAGGGCGAAATTTT
ATTTAAACTCAATCATATGTAAGCGATGAGTTACCTCTCCGCCCCA
CAACAGATCCTTTTAGGATTAGAGAGTAATAT
GTAAAACTTTGAGTGCCCGAAGTATTAGACTTTACTAGAGCC
TGTCATCAATATAACAAAGAAGATAACC
GTCTGACTGTGAAAGGAACAACATTATT
ACCTAATAGAACGTGGACTTTTCCTGAG
TGAGCATAGGCAAATCAACGTAACAAAGCCCTGTACAG
TCCCAAGCGGCTGTTTGATCCTTTAACATTATCATTTTAAAA
TAATGAAGAAATGCAGATGTGCAGCAGACGACAACACTTTAGGAA
GAAACGATTCGTTTTAAGCATCAAATCAGGTCCAG
TGAATAATGGAAAGGAGCAAAAAGCTACC
TTAACAAGAGCCAGAAAGTAATTCCTCA
CCGTTGTAGCGCGAGGCTACGTGGATCGGCAGTATCATAAAGAAA
CCTGCAATCATTCCAAGAAAACATGTAG
TCATCATATTATACGATGCAAGGTTATAATCA
AAGGAATGAGTGAGCGATCTAAAGTTTTCTGTAGCAGGTGAAGCCC
CTGTAATGGAAACAGTAGTTACAATCGGGAGTCAG
TTTTCCCTAAAACATCGCCATTACTATCGGAGAGTCT
CAGTACCGGTCGAGGTGCCTTCTCCAAC
TATCCACCCTATCAAAATATTAGCGTTT
TCGCTATAAGTAAAGCACTAAGAACGTG
GGTTAATAAGAGAATATGAGAATCCAACGCTATCATAACCGA
CAGCAAACGATTCACCCTCATTCGCACT
TTTCATACAGAGCCACCACGACGAGAGGCATTGGCCAATATTACC
GATATTAAAAGCTTGCTTTCGATTCCACCTCATTT
CCAAGAACCGAGCCCGGTCGTCTGCAACAGGAAAAACGCTCATGG
TTTTGGCTTAATTAATTTT
CCAAAACACTTTTGTATCCTGCTCCTGACCAACTT
ATGGCAATGTAACCTATTAAATAAGAATAAAATTATCAGATG
ACTGGATCGTTTACATCCGGTAACGACACAGTTTGGGATTCTCCGT
GAAGTCCGTGACTCTCAATGACGATACGGCGACTTTTGCATAAAG

TTCATTACGGGGTTGCTTTCTGCGCACTGCATCCTGGGCAAC
TTTTTGAACAAGAAAAATAAAACAATACTAATGCCGACGACAAAA
CCGTTTAGTTAGAATCAGCACGTAGCTTAATTGCGCGTCCAATAA
TTTTGACAACTTAAAGGCGCCACCGCGTGTTGCAGCGTGAGTATT
TATTGTGATAAGAGATAGCGAACTGATAGTTTT
GCCAGCCATTGAAATGGATTAT
GAGACGGTACACGCTAAATTTATCCCAATCCCCGC
TCTTCGACCGGAGATCTGGAGCAAACGTCAGATGA
AAAAACAGCTACAAACTCCCAATAGGAACCCTTAGTACCCGCTTT
GAATTGAGTTAAGCAACCACCAGGAAGGAATAGTGTAAT
ACAATCCGCGTATTAGACATCATTGATT
ATTTTGACGCTCAAAATAGGTTAGCGGGGACT
AATCGATCAGAAAATGAATTTCGCGTCGTAAC
TCATCAGTTGAGATATCATAACTGTAGTCGTTTAGTTGTAAAAAAAATA
CATGTGAGTGAGGTCAATGGGCAAAGGA
GTCGAGTCCACCTCCCGTAAGAACAATACTTCTTTGTCATTACCGCTTA
ACTTCAACCAGTTTCTTGTTGTTCGCCACTGGTGAATTGTGCGAGA
TTTTCATGTTCAGGATAAGTCCTTTT
CGCATGAAACATTAGCATATCACCCGGAAATTATTCATCCAG
TTTTGCATTAACAGGTTGTACCAAATTTT
AACATAACTCGTATTAAATGGTGGAAGA
CTGCTACGAATTTCATGCAGCAGCGAAAGACCAGG
CTCAAATATCGAACCATCGCTAACCGTTTATTTTTGATCGGTGCGGGCC
ACAAAGAGGAATTACGAAATCCTCATTAAGTGTAC
TTAGCTTTAAAATACTGCGGAATCTTCGCATTTCA
GCCGTTTGCCCCGTAATATCACCACCATTTGGGGAGGGAAGG
GAGAACCATCTTTCTTAGGAGCCTAATAATTTTTTCACCAAC
GGTAGCAACATTCGGTCGCTGCCTCAGATGCGCTATGTTATCTCGGATG
TATCAGAGAGATAAAACACCCAGCCGGCATCGGAAGCTT
TCCCCATTGGTAACTTTGACGAGTGAGCCTTATCATCCATATAACAGTT
TGTGTTTATAAGGAAGAAGCGCCATTCGCCAGTTG
ACGAAGGACAGTAGGGAAACTCTCAGAATAGC
GGAAATCGTATCATTGCGTGAGCGAGTAACATCTGGCCAGCAAAT
TAACCCTTATTAACGTCGAACCTACCATATCGCGG
ACAGGTATAAAAGATCTTCACCCCCGGTTTGTATATTCCTGTAGCCAGC
CCCTCAGGTTAGTAACCATTGAATCCCC
CGTTGCACCCAGCTACATAGCGAACTAT
AGGTGAGTGTAGGCTATAATTAGCAATCAGA
AATGGTTTTGTTGGGGCATAACCTGTTTAGCTATTTT
TAATTTCATTTGAATTAGAATTATATAGTAA
GAAAAAATTAATTACATTGAGCAATAGGTTGATCCAATAATA
GCGAACGCAATTACCGAAGCCCTTTTTAAGAAAGGAAACTCCTTAATAC
AACTAACATATTGTGAATTACACGAAAGGGCTTTGGAACGAG
TAGAAGAGTCGAAGAAAGCGAAAGGTCA
ATTCGCGACCCGTCAGAAGGCTATCAGGAAACTAG
TCCATCACCAGACGACGTGCGCAACTGTTGGGATGTGCATTTTAACTGT
GTTAAATCAATTCCGAACAAGAGAGTTGCAGGGTATTCACTT
GCACCCAAAAGAACTAAAAGA
AAGACAACGCGTCGTCTCAACAGTTTCAGCGTGCGAATTTAA
CGCTAACGTGTATCAAATAACTATTCATTTCAATT
AACGCATCAGCAAACAGTAGCCCCCTCAGAGCCGCTCACAAA
CCACCAGAGGGTTAAGATGAATATACGTTCTTCTG
TACCACATTCAACTAGGCATAGTACTCATTACATATAAAATTTTAA
TCCGGCTAAGAAGATGATGAA

TAGCAATAGCTATCTAATAAC
ACCAGGCAAGAGGAGTCGAAATATCGGTTTATCTGACAACTA
GGAATGCGAATCAAATCCAGTTACAAAA
CAAATCGCTGAAACATAAGTGCCATTCT
TTTTAACATTATGACCATGCAATGGTGAGAGATA
GAATCTTACCAGCCAGAACCTCGTTAATTTCAAAATGCTTTC
AGGAAGCCAGACTTGAGGTAGCACCAACATATGGCATGATTAAGACCGA
ACCTTAACAATTTTCCCTTAGAATCGCTGAGGTCTGAGTTGA
TTAGGATGTATCACGAGCCGCGCCACCACATTTTCGGTCATAGTAG
CCAATAGGAAAACAAACAACCGTGCGGCCTCAGTC
GCAAACTCCATTCAAAGCGAGTACCTGA
CCTCAGGATAAATTTCGACATGAGTATCTGCATGCTTAAGAG
AGACACCCTCCCAGAGCTCAGACGAGCG
TTTTTATTTTCATTTTTATGGAGAT
GGGCGCCATCCGTTAATATTTTGTAACATTGCAAGGAGTCAA
GGGTCAAAAGGCCTGAGGCTGGCGAAAGGGGGAAG
GGGACGTTTAAAGGGAGCCTAATTTGCT
GTCAAAGCGCATTAGACGGACAAATAAGCCT
CCGAATGGAAATTGGCCAGGCTGAGTTTTGCTCAGTACTAAG
GGATCAGACTGCCCCCTTCACCGGAACC
GGTTGCTAGACTACCTTTTGCAGCTTAGCGCTAGGAGTTACCTCTCCGCCCCA
ATCAATAGCAGAGGCATTTTCCGC
GCCCATTACCCTGGCTGACCTTCAAACCGAACATGTTATCCA
CATCAATAATCATACAGGCAACTCAGAGCGGGAGATAGAACCTCACAAA
GAGTGCTAGCGTCCACAGTTCAGAAAACGAAGAGTTGTC
CAGCATGGCTTTTGATGGGGTCAGAATG
CGCGCGCCGCCCATATTCGAGCGTCTTT
AATAGAACCGCCTGAACCACCGGCGATC
ACGTACACTAACCTAAACTTATGCACGTTAATAAAACG
TTAGTTGATACAGGCGGATGAAAGATTTCGGAATAAGTTTTAACGATAC
CGCCATGAATTCTGCAAAGGATAAAACGCCAGGGTTTTCAGG
ACTGCAATGCGGCACTGTTGACCAAATTCTGCGAACGAGTTTTT
TTGTCCGCGACATCGCCGAATAAGTTTAATTTCAACTT
AAACCTTTAATTGCGGTTCGACAGACCTATTTATTATTTCAATTAAGCA
CCAGATTTTAAGCTAAATTTTTCACGCAAGATCCTGATTGTT
CAACAAATTCTCTGTTTAAAAATTTAATGGTTTGAA
GCCGTCACCAGTTTTCATTACCAGAAAGTAAGCAGATA
GGCCAACAATAAGGCGTTATTAGAAAACCAGGAGGGAACTAT
CTGTAAAGATAAAGAAACCAGAGCCACCACCAGACAGC
AGACACCACGCAACCGATTCATCAAGAAAACAACATAGCGAA
TCAAATATCTTTAGGAGCACTAAAGTCTGGTC
CTAAATCTCCAATATCTACTAATAGTAGTATTTT
GGCGAGAGATCTACAGGTCAATATAAGGCCGTGGA
AGGGGGTAATAACCAAAGGCGTTATTTATAAGCAT
ATAAAGCGGCAAAGATCTGCCACGGTGG
TTTTGCAGAGCAGGCTGGATTATCAAGGCAG
CCCCCTGCCTTATTAAGTTGATATCACCAGAAGCAGCATTTA
TTACGAGAACAGAGGTGAGCTACCACCAGCCTGAGTTAA
ATGGTCAGCGAGCTGAAAATCAACTCGCCTGGGGAACCATCGCCGAAAG
CAGCTCATATTGCAAGGCGATTAATTCAGGCACAGTATCATC
GCTGCTGGCAGTCATAAATATTGGCTACAGAAGATGAT
TTCGCCATGTGCGACATGAAGTTATGTAAAGA
TGGTTTTACAAATCGCGCAGAGGCCCTTTTTTAAATCG
ACGAAGACCCCAATACACGACCAGGCGTATTGCTAAACCGTTTT

GATTCCCTTAGATACATTTAAGAAATTAAAGACGG
CCAGAGTCCAGATCGGTTTTGGAAAGATAGTTACCTCTCCGCCCCA
AATATCCAGAAAGATTCACCAGTCACACGAAACCTTGCTGGT
GCGGCTATTTCAAAAATGAAAATAGCAGCTTTT
TTTTTAAATCAGCAATGGGATAGGTTTTT
TTTTAAATAAAGAATAGGGTGAAAATCTCCA
TTTTTACGTGAACCATAACGGATTCGCCTTTT
TTTTTTATCTTACTCAATGAGTTAGTTTT
TTTTAGTGTTGTTTTGAAGCTAGAAGGCGCCCAAATTT
TTTTTAAAGTACGGTGTCTGGAAGTTTCATGTAAACATGCC
GGTCATTATAATGCTGTACAAAACATATATTATCACGAACCAGATTTT
AGCGCAAAGATGGCAACCAACGGCCAGTCGTGCAAAATTTT
AAACCAATCAATAATCGGCTTTT
TTTTGTCAGAAGCAAAGTATAGATATAGTTTT
TTTTCCAGCAGGCTGTTTT
TACGCCAAGCACGACGTTGTAAAACGTTTT
CGCTGATTGCCGCACTCATCGAGAACAAGTTTT
TTTTAATCAGATACTTTTT
GTCAATAATCTAAACAGTTGAAAGGAATTGTTTT
TTTTTGATTGCTTTGAATACCAACATAAATCAATATTTT
CAATCGCACTGATGGGCATGAGTATGTTTCTAACATCGGGTTGAGTATTTT
TTGATAGCGAAATTACGAATGCAGATACATAACTTTT
TTTTCGCTTCTGGTGCCGGAAACCCCAGTCTTTATTA
TTTTTTTAGAGCTTGACGGGGAATGAACAATAAAAACTTAA
CGTGGCGGATTGACCGTTCATTTTCGCATTAAATTTTTGTTTTT
TTTTATTGAGCGCTAA
TTTTGCCAAAAGGGAGGCTTTTGCATTTT
TTTTTATGTGAGTGAATAACCTTCCCTAAAATCAAGTAACC
TTTTAGTCTGAGCAAAGATGAAGGCGGATTTTCGAGCACAGGTCGATAAGA
TTTTCAGCAATAAAGACGCCTTACATA
ATAGGAACAATATCAGGAATAACAAGTCAGAGGGTATTTT
TTATAGATTTAAACAATCACCCAAGGGAGCCCCCGATTTT
TTTTCCATGGCTTAGTTTT
TTTTCTTTACAGAGAGGCGATGGCCCACTTTT
TTTTACGGGCAACAGCTGGTTTGCCTTTT
TTTTTTAGATGATTATTTT
TTTTAAAGAAGTTTTGCTTTACCCTGACTATTATATTTT
TTTTAACCCAATATGTTTT
TTTTGAGGATTTAGAACGTTATTAATTTT
TTTTCAAGCCGTTTTTTAGCAAGCATTTT
TTTTATTAATGAACATTTT
TTTTACAGTTCAGGGTTTT
TTTTAGTCGGCATACAGCTTTCCGGCACTTTT
TTTTTGTCTTTCCTTACAGTGCCAAGCATCAAA
TTTTAGCTTAATTGCTGAATTTTGCGGGGAA
CGATAGAGGACTTGTGCAAGTGATGTTTTAAATATGCAACTTTT
TTTTGTCGTTCAGGGTTGTCTGAACGAAAATGCGGAATTAC
TTTTTTTTAAAAGAGTTTT
TTTTAGGAAGGTTGATAATACATTTTTTT
TTTTCACGTTGGTGTACCAGCCAAATATTCGAAATGAGGTG
TTTTTAAATCAAGATTAGTTGGAGGTTCCAGTTTGCCCGAGAATTGCGTTTGCAC

#### Seam staples

Every line describes the sequence of one staple.

ATTAAACGGGCCGGAACCGGATCGTCACCCTAGGAAGTTTCC
TCTATGGTTTATAAGACGGTCAATCATAAGGGTCAAGAGTAA
TTTTTCCAGACGTTAGTCACCAGTTGATACC
TTTTCGCCCGTATACAAGAGAAGGCAGGCGCC
AAATAAATCAAAACTCAACGAGAGCTTG
GCAGTCAGGACGTTGGGAAGAAAAATCTGATTTTAGAGATGGGCTT
AATAGGCCGGAAAATACTTACGCAGTATGTTAACAAAGTCGGTAA
TTGAGACTGAGTTTCGTAAATGAA
CGAAACAGGAACGAGGCGCAAATTTACCGAGG
TCTCCAGTAAGGTTGTATGGGATTTTGCTAAAGTTGAAAATC
CGCATCAATAGAAAATTAACGTAGAAAAGCG
AAACAGCAGGAGGTATGTACCGTAACGA
GATAGAGGTCAAAAAAAAGGCCTTAGCCAAGTACAGAAACAC
AATACGTAGCGATTATACCACAGCATTGTCCCTCACGTACTCAACCTAC
ATACCCGAGCGAGTAGTAAATTGGGCTTAGAA
TGCGGCCACTTGCGCCGACAAGACCCCCAATGCCAGCTCATT
CAGAACACATTGACAAGAACCGGATATTCTGACGAACGGAGACATC

#### Force staples

Every line describes the sequence of one staple.

### 15.6 pN

GAGCTAAAAATCTAACGCTGAGAGCCAGCATTTT
ATTAAACAAATTAGTAATAACATCACTTGCAGAAGGACAATAAGTGTAAGTT
ATACTTAGTCTACATACGATAAAGCGCCATAT
AGTAGCCGGAGTCCAGAAGAACGCGCGCGGT
TTTTGCAAATGAACTCACATTAATTTT
GGTGCCTCGCCTGGCCCTGCAATCGGGT
ATACATGTAATTTCAACTAATAGATAAACAATTCGACGCAGCCTGG
TTTTAGTACCGACAATAAACCCACACATTAATGCGAACCCTCACC
TGGATTCCTGCACCGGACAACAGTCACAATTAATTTT
TCATCGTAGGCAAGTACCCTTCACAATGAGT
CAGTATTCAAATATCAAACCTAGCATAATTTTTGAGTAAGAAGTTTTTT

### 29.6 pN

GGGGTGCAAATCTAACGCTGAGAGCCAGCATTTT
ATTAAACAAATTAGTAATAACATCACTTGCAGAAGGACAATACGAGCCGGTT
ATACTTAGTCTCTCACAAATAAAGCGCCATAT
AGTTTCCACAGTCCAGAAGAACGCGCGCGGT
TTTTGCAAATGAACTAATGAGTGAGTT
AAAGTGTCGCCTGGCCCTGCAATCGGGT
ATACATGTAATTTCAACTAATAGATAAACAATTCGACGCGAAGCAT
TTTTAGTACCGACAATAAACTTATCCGTTAATGCGAACCCTCACC
TGGATTCCTGCACCGGACAACAGTGAAATTGAATTTT
TCATCGTAGGCAAGTACCCTTCACAAAGCCT

CAGTATTCAAATATCAAACCTCAACATATTTTTGAGTAAGAAGTTTTTT

**35.2 pN**

GCCTGGGAAATCTAACGCTGAGAGCCAGCATTTT
ATTAAACAAATTAGTAATAACATCACTTGCAGAAGGACAATACATACGAGTT
ATACTTAGTCTTCCGCTCATAAAGCGCCATAT
AGTACAATTCGTCCAGAAGAACGCGCGCGGT
TTTTGCAAATGAAGTGCCTAATGAGTT
GCATAAACGCCTGGCCCTGCAATCGGGT
ATACATGTAATTTCAACTAATAGATAAACAATTCGACGCGCCGGAA
TTTTAGTACCGACAATAAACATTGTTATTAATGCGAACCCTCACC
TGGATTCCTGCACCGGACAACAGTGTGTGAAAATTTT
TCATCGTAGGCAAGTACCCTTCACGTGTAAA
CAGTATTCAAATATCAAACCTCACACAATTTTTGAGTAAGAAGTTTTTT

**38.8 pN**

AAGCCTGAAATCTAACGCTGAGAGCCAGCATTTT
ATTAAACAAATTAGTAATAACATCACTTGCAGAAGGACAATAAACATACGTT
ATACTTAGTCTTATCCGCATAAAGCGCCATAT
AGTTCACAATGTCCAGAAGAACGCGCGCGGT
TTTTGCAAATGAAGGGTGCCTAATGTT
AAGCATACGCCTGGCCCTGCAATCGGGT
ATACATGTAATTTCAACTAATAGATAAACAATTCGACGCGAGCCGG
TTTTAGTACCGACAATAAACAAATTGTTTAATGCGAACCCTCACC
TGGATTCCTGCACCGGACAACAGTCTGTGTGAATTTT
TCATCGTAGGCAAGTACCCTTCACAAGTGTA
CAGTATTCAAATATCAAACCTTCCACACTTTTTGAGTAAGAAGTTTTTT

#### Probing staples

Probing staple for EcoRV experiments:

5' - Cy5 - GGAGATATCAATTCGTAATCATGGTCAT - ATTO 550 - 3'

Probing staple for EcoRI experiments:

5' - ATTO 550 - ATCCCCGGGTACCGAGCTCGAATTCGTA - Cy5 - 3'

Probing staple for BamHI experiments:

5' - Cy5 - AATGGATCCCCGGGTACCGAGCTCGAAT - ATTO 550 - 3'
